# ComBatLS: A location- and scale-preserving method for multi-site image harmonization

**DOI:** 10.1101/2024.06.21.599875

**Authors:** Margaret Gardner, Russell T. Shinohara, Richard A.I. Bethlehem, Rafael Romero-Garcia, Varun Warrier, Lena Dorfschmidt, Lifespan Brain Chart Consortium, Sheila Shanmugan, Paul Thompson, Jakob Seidlitz, Aaron F. Alexander-Bloch, Andrew A. Chen

## Abstract

Recent work has leveraged massive datasets and advanced harmonization methods to construct normative models of neuroanatomical features and benchmark individuals’ morphology. However, current harmonization tools do not preserve the effects of biological covariates including sex and age on features’ variances; this failure may induce error in normative scores, particularly when such factors are distributed unequally across sites. Here, we introduce a new extension of the popular ComBat harmonization method, ComBatLS, that preserves biological variance in features’ locations and scales. We use UK Biobank data to show that ComBatLS robustly replicates individuals’ normative scores better than other ComBat methods when subjects are assigned to sex-imbalanced synthetic “sites”. Additionally, we demonstrate that ComBatLS significantly reduces sex biases in normative scores compared to traditional methods. Finally, we show that ComBatLS successfully harmonizes consortium data collected across over 50 studies. R implementation of ComBatLS is available at https://github.com/andy1764/ComBatFamily.

## Introduction

Neuroimaging research’s recent shift towards large sample sizes necessitates pooling data collected across many sites and scanners^1–3^. Both large consortia and multisite studies often use statistical methods to correct batch effects resulting from subtle differences in MRI acquisition, hardware, or collection protocols across sites^4,5^. One popular technique is ComBat^6^, which estimates and removes site-specific offsets in feature distributions while preserving the linear effects of prespecified covariates^7,8^. This framework has been extended to create several new methods, including ComBat-GAM, which preserves nonlinear covariate effects via generalized additive models^9^. Collectively, these methods perform well in large samples across a range of imaging modalities^10,11^ and increase statistical power^12,13^. However, while they preserve covariate effects on features’ means, no existing harmonization methods preserve the effects of biological covariates on neuroanatomical features’ variances.

These biological sources of variance are integral to brain phenotypes’ distributions across a population. Recent literature has demonstrated that factors such as age and biological sex impact the variances of neuroanatomical features^14–18^. While mean differences have received greater attention, sex differences in brain structures’ inter-individual variabilities are common and may impact disease prevalence^14–16,19^. Similarly, assessments in adolescents^18^, elderly populations^17^, and across the lifespan^20,21^ have found age-related differences in the variances of several neuroanatomical features.

Biologically valid models of features’ scales, or distributional dispersions, and robust harmonization are critical for generating normative trajectories of neuroanatomical features. This longstanding goal of neuroimaging research has recently become feasible thanks to massive consortia and open-access datasets^20–23^. Besides mapping a normative trajectory of brain structure throughout life, these models make it possible to quantify each individual’s offset from this norm as a percentile (centile) or Z-score. Crucially, these scores depend on accurate models of population-level variance relative to factors like age and sex, making them highly susceptible to the loss of biological variability imposed by current harmonization methods.

Existing techniques’ failure to preserve biological scale effects may be particularly problematic when such variance-altering phenotypes are distributed unequally across sites^24^. For example, current ComBat methods would force sites with male-dominated and female-dominated samples to target the same variance despite known sex differences in features’ scales^8^. While this may be avoided by harmonizing males and females separately, this comes at the expense of statistical power and is not possible for continuous covariates like age. Thus, by removing biological effects on feature variance, harmonization may produce inaccurate or biased estimates of normative scores, which in turn could yield inaccurate or biased associations with phenotypes such as clinical diagnosis. Therefore, preserving the effects of covariates like age and sex on feature variance is crucial to estimating accurate normative scores for all individuals.

In this work, we extend the ComBat framework to perform batch correction while flexibly preserving covariates’ effects on each feature’s location and scale. This new method, ComBatLS, integrates generalized additive models for location, scale, and shape (GAMLSS)^25^ to preserve complex, nonlinear effects in both the first and second order of a feature’s distribution during harmonization. We first assess ComBatLS’s ability to preserve covariate effects on scale by applying it to synthetic “sites” containing unequal numbers of males and females, which we created from UK Biobank data^26^. Here we focus on sex since it is commonly imbalanced in biomedical research samples^27–29^ and its effects on neuroanatomical variance are well-documented, including in this sample^15^. We hypothesized that normative scores calculated from data harmonized across synthetic sites with ComBatLS would recapitulate subjects’ true scores more accurately than those derived from data harmonized with other ComBat methods. We also conducted several tests for sex biases in how each method impacts score estimates, hypothesizing that ComBatLS would reduce sex-related biases more effectively than other ComBat methods. Finally, we validated that ComBatLS successfully harmonizes massive datasets with varying demographics by applying it to data from the Lifespan Brain Chart Consortium (LBCC)^20^ collected by over 50 primary studies. We have released ComBatLS as part of the ComBat family of R methods at https://github.com/andy1764/ComBatFamily.

## Results

### The ComBatLS method for multi-site harmonization

CombatLS extends the original ComBat framework by flexibly modeling and preserving the effects of specified covariates on each feature’s location and scale. The effect of selected covariates is fit using a generalized additive model for location, scale, and shape (GAMLSS)^25^. This framework directly models the first and second moments of each feature’s distribution before shrinking site parameter estimates towards their mean estimate across features, enabling ComBatLS to preserve non-linear covariate effects in both mean and variance while removing batch effects.

To assess ComBatLS’s ability to preserve the biologically-derived variance necessary for calculating centile scores, we randomly sampled data from the UK Biobank (N=28,619, 49.7% female) to create three synthetic sites, two with unequal sex ratios (**Figure 1**). We used ComBatLS to “harmonize” neuroanatomical features across these synthetic sites while preserving the effects of age and sex. Then, we compared subjects’ centile scores from brain charts fit on these data to true centile scores derived from the unharmonized dataset. Since centile scores are defined by the population’s distribution, recapitulating these true centile scores requires that harmonization preserve any between-site differences in variance resulting from the samples’ imbalanced sex compositions. We quantified the offset of each subject’s ComBat-derived centile scores from their true centiles as “centile error”, with smaller errors indicating better variance preservation. We then compared the distributions of centile errors and their magnitudes when data were harmonized by ComBatLS and three other methods: linear ComBat, ComBat-GAM, and ComBat without any covariate preservation. Results from ComBat without any covariate preservation – a theoretical “lower bound” to harmonization which performed worst across all analyses – are presented in the Supplement (Section I, SFig 1-10). To account for the randomness in creating synthetic sites, we resampled subjects’ site assignments and repeated these analyses 100 times, which produced highly consistent results across replications (Supplement Section I).

**Figure 1.**
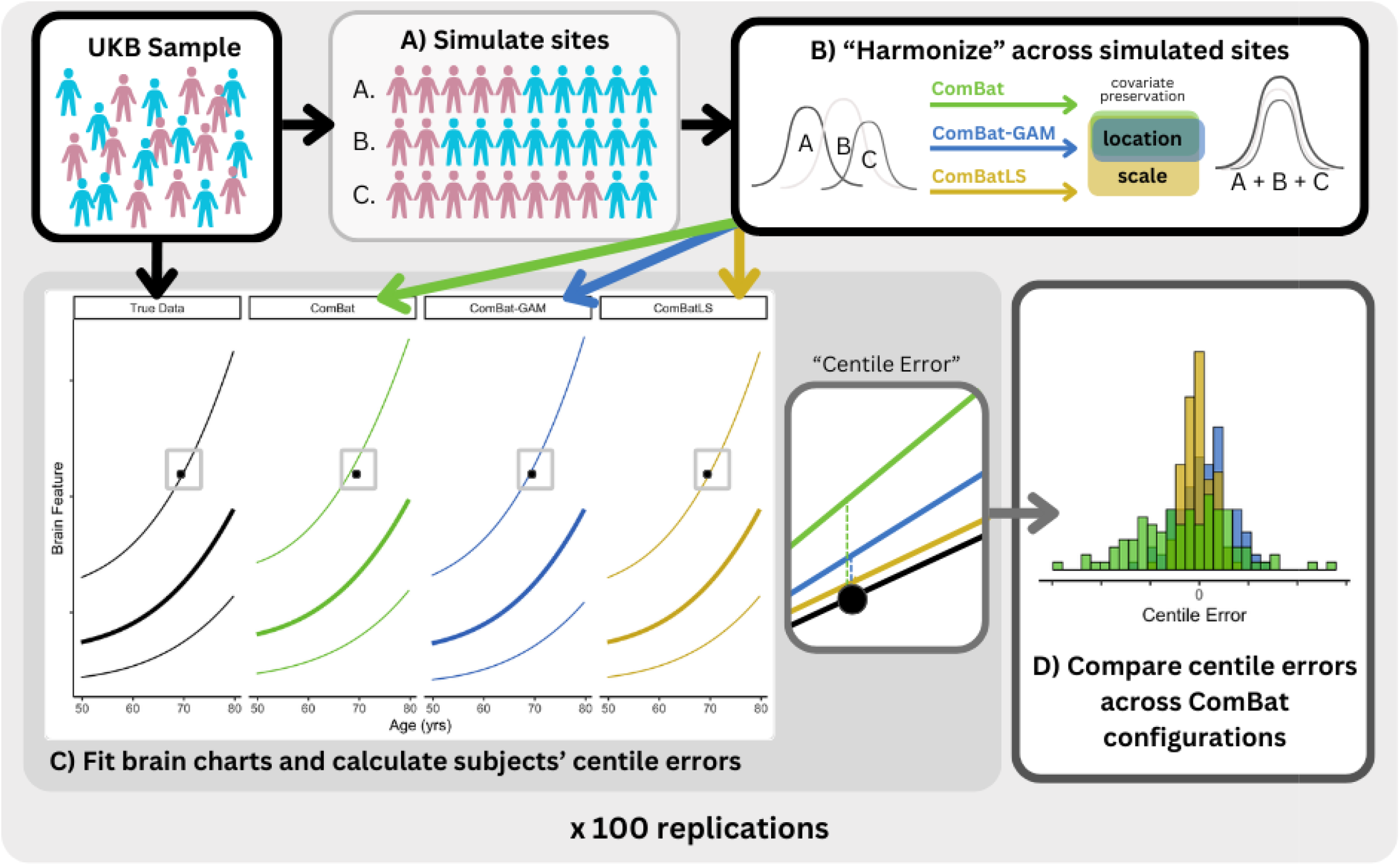
Summary of Methodological Approach. A)Subjects from the UK Biobank (UKB) sample are randomly assigned to one of 3 simulated “study sites” such that sites have a Male:Female ratio of 1:1, 1:4, or 4:1. B) Brain structure feature data are “harmonized” across these simulated sites using different configurations of ComBat, preserving covariates’ effects on the mean (ComBatLS, ComBat-GAM, linear ComBat) and/or variance (ComBatLS). C) Brain growth charts are fit for brain features using the original true structural data and data harmonized by each ComBat pipeline, which are then used to calculate personalized centile scores describing each subject’s percentile relative to the population distribution. Centile error, defined as the difference between a subject’s centile score when benchmarked on a brain chart modeled on “true” data and one fit on ComBat-harmonized data, was calculated for each brain feature across all subjects. Lines represent brain charts of the 75th, 50th, and 25th percentiles for the feature given age; the solid point represents a single subject’s brain feature, which has a “true” centile of 75% but corresponds to different centile scores when data is harmonized. D) We analyzed the distributions of centile errors within and between ComBat configurations to assess the degree to which each harmonization method preserved biological variability in the simulated sites, thus minimizing centile errors.

### ComBatLS recapitulates centile scores across sex-imbalanced sites better than other ComBat methods

We first fit GAMLSS models to derive centile scores from 208 neuroanatomical features – four global tissue volumes and 68 cortical regions’ volume, surface area, and thickness measures – harmonized by each ComBat method (**Figure 2A**, see Methods). Using non-parametric, paired, two-sample tests of absolute centile errors, we found that in nearly all features, ComBatLS had significantly lower absolute centile errors than linear ComBat or ComBat-GAM (**Figure 2B-C;** Supplement Section I: Table 1, SFig. 1-3), indicating that this method induces less error in centile scores when harmonizing across sex-imbalanced simulated sites. Across our 100 replications, all volume and surface area features’ absolute centile errors were lowest when fit with ComBatLS. However, a small number of cortical thickness features were fit better by ComBat-GAM or ComBat (SFig. 2), which both performed better in cortical thickness than other features. This is expected because, as in prior studies^14–16,30^, the variance of cortical thickness features is less impacted by sex than the variances of other features modeled in this sample (Supplement Section II, SFig. 11); therefore less information is lost when harmonization does not preserve these effects. ComBatLS’s added utility beyond other ComBat methods is thus proportional to the degree to which preserved covariates impact features’ variances, though it performs well even in the absence of substantial scale effects. In sum, these results demonstrate that ComBatLS is highly effective in preserving biological covariates’ effects on features’ scales, even when those covariates are not distributed evenly across study sites.

**Figure 2.**
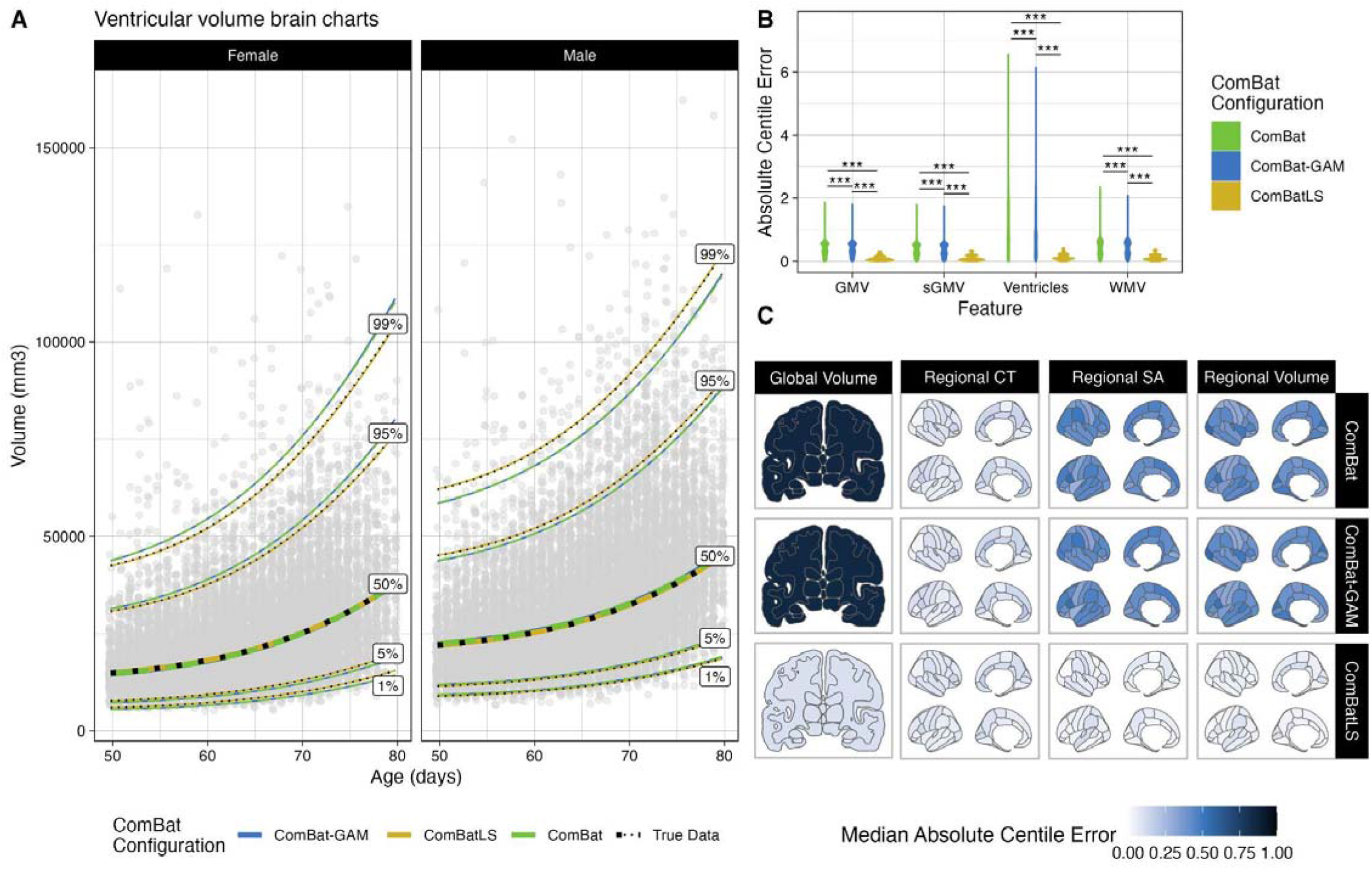
ComBatLS best replicates centile predictions across simulated sex-imbalanced sites. A) Brain charts of ventricular volume in females and males when derived from data harmonized across simulated sites with ComBat, ComBat-GAM, or ComBatLS relative to unharmonized (“true”) data. Line color corresponds to ComBat configuration, black dotted line corresponds to true data. Age values jittered slightly for visualization. Absolute centile errors, or distance in centile space between subjects’ true centiles and centiles derived from data harmonized with different ComBat configurations, of global volumetric features. C) Median absolute centile errors across brain features in data harmonized with varying ComBat configurations. Abbrv: WMV, white matter volume, GMV, cortical gray matter volume, sGMV, subcortical gray matter volume, CT, cortical thickness; SA, surface area; ***, p < 0.001, FDR-corrected.

In addition to replicating these analyses after re-sampling subjects’ site assignment 100 times (Supplemental Table 1), we assessed whether extreme phenotypes drove the observed results by removing centile scores >95% or <5% when calculated on the original data. Again, these results were highly similar to those of the full dataset (Supplement Section III, SFig. 12-14). We also repeated these analyses using Z-scores (theoretical range (-inf,inf)) which yielded a nearly identical pattern of results to centile scores (range (0,1))(Supplement Section IV, SFig. 15-25). Together, these analyses indicate that ComBatLS is consistent and robust to extreme phenotypes and choice of deviation score.

### Sex differences induced by ComBatLS are small and less directionally biased than other ComBat methods

We next assessed whether the centile errors induced by each ComBat method vary by subject sex, producing centile estimates that could bias associations of interest. We first mapped significant effects of sex on variance estimated by each feature’s brain charts, which revealed that models fit on ComBatLS-harmonized synthetic data closely match effects quantified in the true data (**Figure 3A**). To see whether these discrepancies in sex estimates affect centile scores, we used Wilcoxon tests to compare males’ and females’ centile errors across ComBat methods within each feature. For all methods, most of the 208 features exhibited significant sex differences in centile errors (ComBatLS: 199 features, ComBat-GAM: 140 features, ComBat: 138 features, FDR-corrected across features and methods; Supplemental Figure 5). Next, to test whether these differences produced sex biases by consistently over- or under-estimating either sex’s centiles, we examined the distributions of these sex-differences’ medians across 100 sampling replications. This revealed that linear ComBat and ComBat-GAM tend to produce slightly more negative sex differences in centile errors, corresponding to a systematic underestimation of males’ centile scores relative to females’ (**Figure 3B**; Supplemental Figures 6 and 7). ComBatLS produced a very small but significant negative male bias only in global tissue volume centiles (β = -3.1e-04 centiles, p=0.0075, FDR-corrected across method and feature categories), while again, cortical thickness was exceptional in that significant biases were also absent in ComBat-GAM’s and ComBat (ComBat-GAM: β = -9.13e-05 centiles, p=0.52, ComBat: β = -9.57e-05 centiles, p=0.48, FDR-corrected). We further visualized whether these biases resulted in over-representation of either sex among subjects with very high (>80%) or very low (<20%) average centiles in any feature category. Interestingly, we found that while ComBatLS, ComBat-GAM and ComBat all tend to slightly over- or under-represent females among those with extreme average centiles, ComBatLS showed less bias compared to other harmonization methods (**Figure 3C**). Together, these results suggest that ComBatLS-harmonized data preserves sex’s effects on variance and thus induces the smallest and least-biasing differences in males’ and females’ centiles.

**Figure 3.**
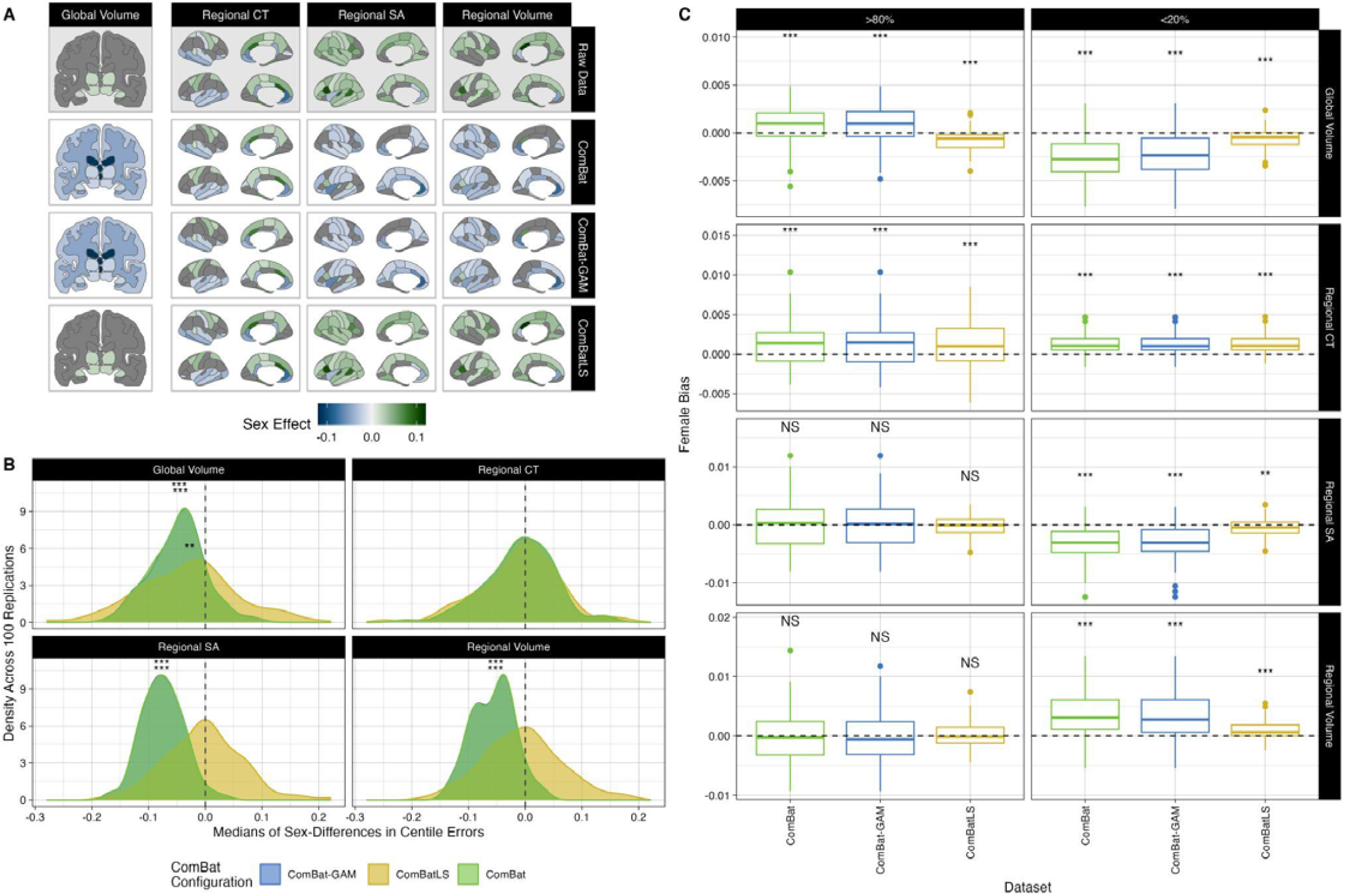
ComBatLS induces less sex biases in centile scores than other ComBat methods. A) Brain features with significant sex effects in scale, as determined by the second moment of GAMLSS growth charts. Fill represents the difference in males’ and females’ predicted variance at the sample’s mean age (64.94 years), standardized by dividing by females’ predicted variance. Positive effects indicating that males’ variance is higher than females’. Gray regions indicate non-significant sex effects. B) Density plots of median sex differences in centile errors induced by different ComBat methods within phenotype categories across 100 replications. C) Bias in the proportion of females with low (<20th percentile) or high (>80th) mean centiles across 100 sampling replications. Positive values indicate a higher proportion of females than “true” mean centiles calculated from unharmonized data (dashed line). Abbrv: CT, cortical thickness; SA, surface area; ***, p < 0.001; **, p < 0.01, FDR-corrected two-sided, one-sample t-tests.

### ComBatLS preserves covariate effects across a wide range of sex imbalances in synthetic sites

To test how each ComBat method’s performance is affected by varying degrees of covariate imbalances, we again drew eleven samplings of UK Biobank participants to create two synthetic sites: one with an even sex ratio and one in which the percentage of males ranged from 0% to 100% in 10% increments (**Figure 4A**). We then repeated the same procedures as in our main analyses to obtain and compare centile errors between ComBat methods when harmonizing each of these eleven samples.

**Figure 4.**
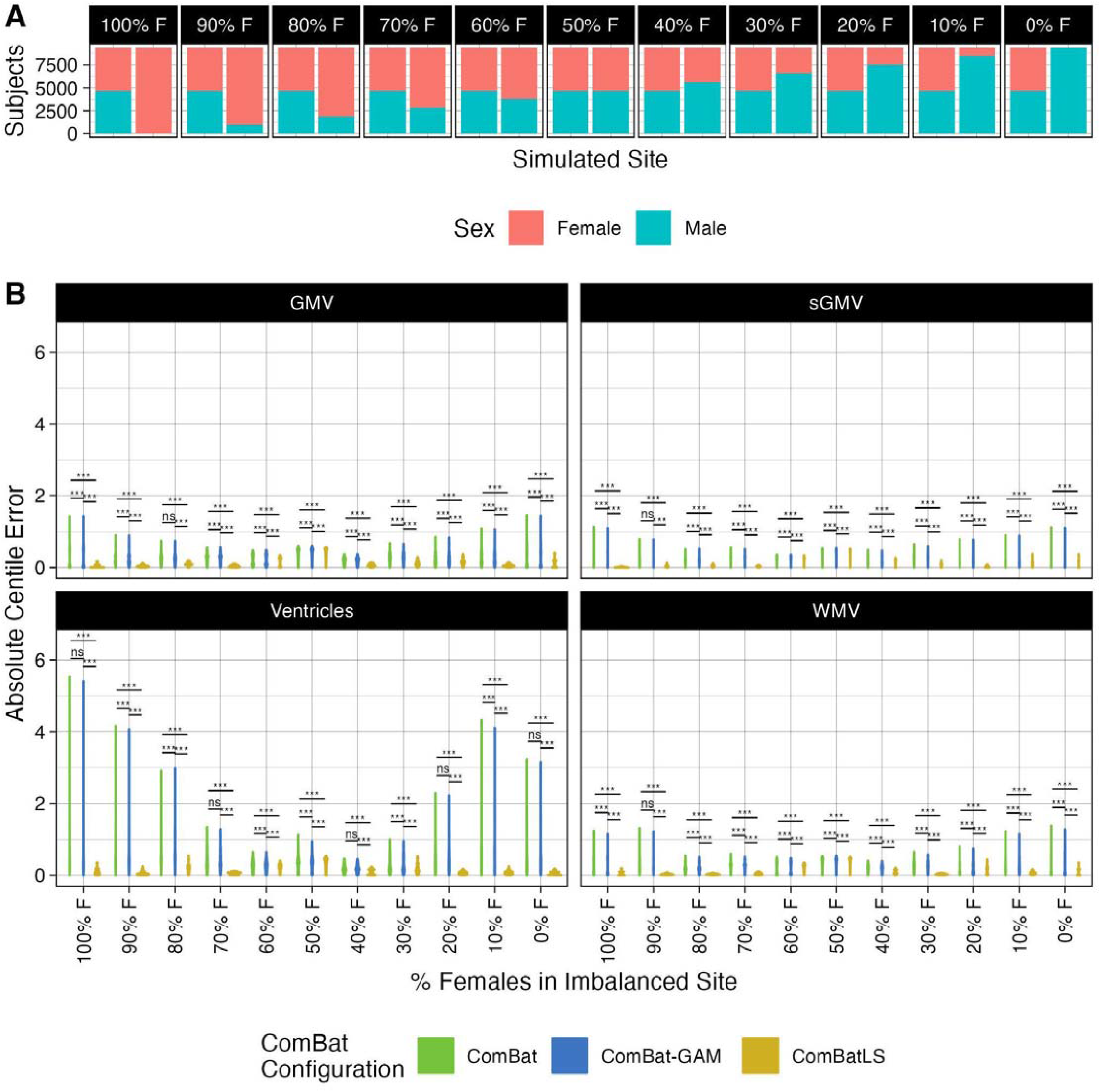
ComBatLS best preserves the effects of biological sex on centile scores across varying degrees of site-level sex imbalance. A) Distribution of Female and Male UKB subjects across two simulated sites, permuted 11 times to assess the performance of ComBat configurations across varying degrees of site-level sex imbalance. B) Distributions of absolute centile errors of global brain features harmonized across sites simulated with varying degrees of sex imbalance using ComBat, ComBat-GAM, and ComBatLS. Abbrv: WMV, white matter volume, GMV, cortical gray matter volume, sGMV, subcortical gray matter volume, CT, cortical thickness; SA, surface area; ***, p < 0.001; **, p < 0.01, FDR-corrected.

**Figure 5.**
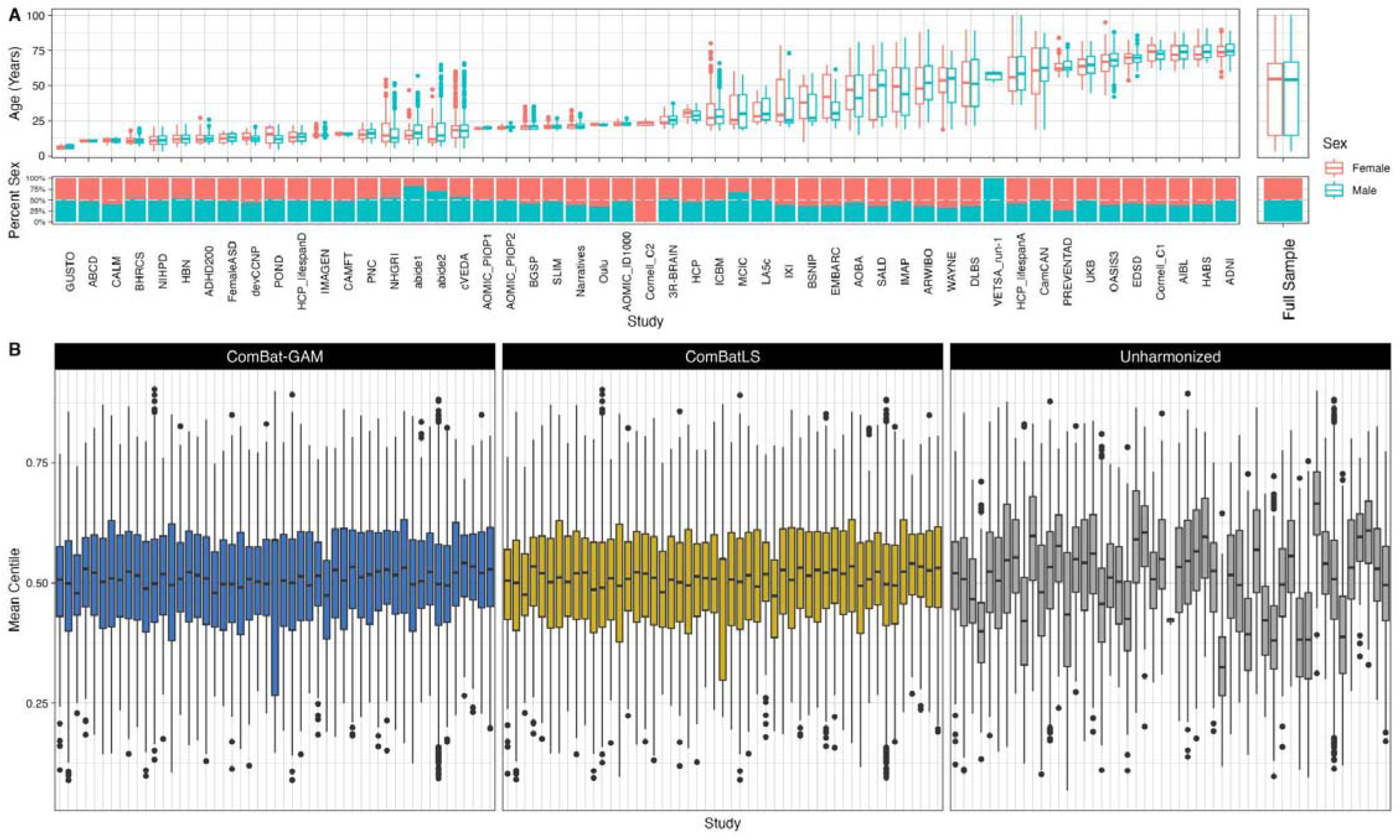
ComBatLS and ComBat-GAM both effectively remove batch effects from consortium data containing over 50 studies. A) Characteristics of curated LBCC sample by primary study and summarized across the full sample. B) Distributions of subjects’ mean centiles across all brain features within each primary study. Centiles are calculated from brain charts fit on data harmonized using ComBat-GAM or ComBatLS, or on unharmonized data, and averaged within subjects across features.

For most features, ComBatLS produced significantly smaller absolute centile errors than other ComBat methods when one site was imbalanced for sex (**Figure 4B**). As above, these gains are less apparent in cortical thickness measures and when both sites are perfectly balanced for sex (Supplemental Figure 9). Our assessment of sex biases in each method’s centile errors was again consistent with our main analyses (Supplemental Figure 10; Supplement Section I).

These analyses demonstrate that ComBatLS adaptively preserves biological covariates’ effects on variance in a range of simulations and becomes more beneficial for ensuring centile accuracy as samples’ compositions become increasingly imbalanced.

### ComBatLS effectively harmonizes massive real-world datasets

To assess its utility in real-world samples, we applied ComBatLS and ComBat-GAM to data from the Lifespan Brain Chart Consortium, a global sample of structural MRI from healthy individuals across the human lifespan^20^. Here we used scans from 52,098 (51.2% female) unique subjects aged 3.2 to 99.9 years collected by 51 primary studies over 199 sites (**Figure 5A**, see Methods for data curation). Both ComBatLS and ComBat-GAM mitigated batch effects across all features as indicated by comparing residual study effects to those in unharmonized data (Cohen’s F-squared: ComBatLS median=0.0087, IQR=0.030; ComBat-GAM median=0.0092, IQR=0.030; Unharmonized median = 0.083, IQR = 0.090; Supplemental Figure 26). However, there are differences in the resulting centile scores when the data is harmonized with ComBatLS or ComBat-GAM (mean absolute difference in centile scores = 0.456, range = 0.032 - 8.42 centiles) with 46.9% of subjects having discrepant categorization of extremely high (< 5%) or low (> 95%) centiles in at least one feature (mean=0.91 features per subject, range = 0 - 49 features). We conducted exploratory analyses to assess whether mean differences in subjects’ ComBatLS- and ComBat-GAM-harmonized centile scores varied as a function of the primary studies’ characteristics, revealing a subtle association with the mean age of the sample and how much an individuals’ age deviated from that sample mean (Supplement Section V, SFigs 26-28). The simulation studies described above (Figures 1-4) suggest that ComBatLS’s accurate preservation of scale effects, including age effects, contributes to the observed differences in centile scores from consortium data harmonized with ComBatLS compared to ComBat-GAM.

## Discussion

Here we introduce and validate ComBatLS, a novel location- and scale-preserving method to harmonize data across batches. Neuroimaging has begun to answer long-standing questions about the brain’s normative structure across the lifespan and to quantify individuals’ deviation from this norm^20–22,31^. However, such studies require integrating vast quantities of MRI scans collected across studies, sites, and scanners, which are highly unlikely to have similar sample characteristics. While ComBat and its extensions preserve the effects of such biological covariates on features’ means^8,9^, failure to preserve variance effects during harmonization may increase error and induce biases in downstream analyses. This includes normative scores derived from growth charts and estimates from brain-age models, which are known to be impacted by harmonization^32,33^.

Furthermore, while additional work is needed to quantify ComBatLS’s impact outside of normative modeling, the relationship between variance and effect size implies that errors in scale could impact group-level inferences^34^. By integrating scale preservation in a robust batch-correction framework and illustrating its ability to improve individuals’ normative scores, this study and corresponding open software represent a significant advance in neuroimaging harmonization.

Leveraging structural features derived from nearly thirty thousand UK Biobank participants, we found that ComBatLS recapitulates subjects’ true normative scores better than ComBat or ComBat-GAM. We first used weighted sampling to create synthetic batches that were male-dominated, female-dominated, or balanced for sex. We then applied ComBatLS and several existing ComBat methods to “harmonize” these data and assessed how each method offset subjects’ centile and Z-scores from their true values. Across 100 sampling replications and 208 brain features, we found that ComBatLS consistently produces normative scores closest to the ground-truth, unharmonized data. This result remained stable across a range of sensitivity analyses, including removing subjects at the extreme ends of the normative distribution and when synthetic samples’ sex differences were as small as 10%. These results indicate that by preserving the effects of biological covariates on distributions’ scales, ComBatLS enables highly accurate quantifications of individuals’ deviations from neuroanatomical norms.

ComBatLS’s preservation of this critical intersubject variability is demonstrated by its performance in features with variances that are both strongly and minimally affected by sex. Across our analyses, harmonization methods were most comparable when applied to cortical thickness features, whose variances are less impacted by sex than other neuroanatomical phenotypes^14–16,30^ (Supplement Section II). Yet, even in cortical thickness, we found that ComBatLS still recapitulates each feature’s true centiles better than the other methods in more than half of replications. Thus, the relationship between ComBatLS’s performance and the strength of sex effects on scale only emphasizes how important such effects are for centile estimation and should not discourage ComBatLS’s use in features with minimal biological impacts on variance.

One motivation in developing ComBatLS was to prevent harmonization from inducing sex biases in normative scores by systematically inflating or deflating the centiles of either sex. Interestingly, our analyses reveal that none of the covariate-preserving methods – ComBat, ComBat-GAM, and ComBatLS – induce large differences in males’ and females’ centile scores (Supplement Section I). However, ComBatLS induces less bias in centile errors across 100 replications, suggesting that covariate preservation in scale does mitigate harmonization-induced biases.

We also validated that ComBatLS can harmonize neuroanatomical data across more than 50 real-world studies with heterogeneous sample demographics, clearly demonstrating the practical scalability of ComBatLS to large, heterogeneous studies. Exploratory assessment of differences in the centiles calculated from ComBatLS or ComBat-GAM, which was developed to harmonize studies spanning wide age ranges^9^, reveals that subjects’ mean absolute differences increase with the mean age of their primary study’s sample and when that mean age is further from their own. Alongside our simulation analyses, these results suggest that by preserving the impact of age on variance, ComBatLS may produce more accurate centile score estimates than ComBat-GAM in real-world datasets.

This work has several limitations. First, as with all ComBat methods, ComBatLS will only preserve covariates that have been pre-specified. Similarly, ComBat methods rely on some degree of between-batch variation to accurately estimate covariate effects, as illustrated by the slight increase in absolute centile errors for all methods when our simulated sites were balanced for sex (see Figure 4B). Thus, while even small sampling imbalances seem sufficient for ComBatLS, researchers should take care when preserving covariates to avoid overfitting. Additionally, ComBatLS does not preserve covariate effects in or remove batch effects from higher-order moments such as skew or kurtosis, which will necessitate new methods to resolve^35^. Finally, while popular, the existing cross-sectional ComBat methods assessed here only represent one flavor of multi-site harmonization^4,5,36,37^. It will be important for future work to compare ComBatLS to – or potentially integrate it with – other innovative and emerging frameworks^38–40^.

The current study introduces ComBatLS, an R-based harmonization tool that extends the ComBat framework by preserving the effects of biological covariates on both location and scale. Its ability to robustly estimate individuals’ normative scores may enable sensitive identification of disease effects, increasing the power of future clinical research^20^. Furthermore, widespread sex differences in the variances of measures beyond neuroanatomy^41–43^ indicate that ComBatLS may be useful for correcting batch effects across a broad range of disciplines. ComBatLS is openly available at https://github.com/andy1764/ComBatFamily.

## Methods

### ComBatLS

ComBatLS is an extension of ComBat-GAM^9^ that leverages the GAMLSS framework to model and preserve covariate’s effects in feature’s locations and scales. Specifically, ComBat-GAM assumes that for site **i**, subject **j**, and feature **k**, *e* _*ijk*_ ∼*N*(0, σ^2^ _*k*_) in

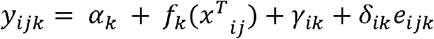

In ComBatLS, we modify our model to incorporate a log-linear relationship^44^ between the error standard deviation and the covariates

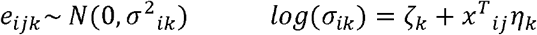

Denote the standard data as 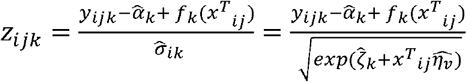. We then obtain our harmonized observations as

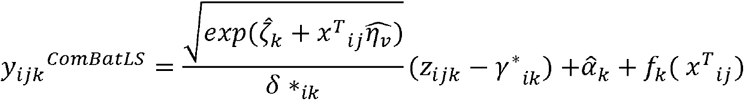

An important consideration arises in how to regress out site during the standardization step. Johnson, Li and Rabinovic^6^ obtain least-squares estimates 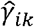 by constraining 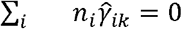 for all *k* 1, 2, …, *p*. ComBatLS, the log-linear model of the error’s standard deviation and covariates is estimated from the site-influenced errors δ_*ik*_ *e*_*ijk*_∼ *N*(0, δ^2^_*ik*_ σ^2^_*ik*_), which implies

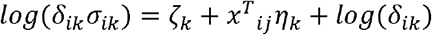

For this to be identifiable^6,8^, we first fit the model without an intercept to obtain estimates <u>*η*_*k*_</u> and <u>*σ*_*k*_</u>. Then, we estimate the intercept as the pooled mean 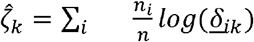. Finally, our estimates 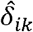 are obtained as deviations from the estimated grand means 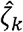 via 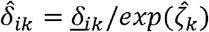. Importantly, ComBatLS retains the strengths of prior ComBat iterations, including empirical Bayes methods to improve batch effect estimates for small sites^6,8^. Similarly, users can interrogate models’ diagnostic plots using the plot.comfam() wrapper. See Hu et al^5^ for recommendations on how to assess harmonization performance.

### Software

All code used to perform analyses and create figures is available at https://github.com/BGDlab/combat-biovar. ComBatLS can be found at https://github.com/andy1764/ComBatFamily. Subject sampling, main analyses, and visualizations were done in R v 4.1.1. ComBat harmonization, sampling, and analyses for replications were done in R v 4.1.2.

### Simulation experiments

#### UK Biobank Data

Our analyses leveraged structural MRI from the UK Biobank (UKB), a large, deeply-phenotyped population sample of adults. Details of neuroimaging data acquisition are available elsewhere^26^. Importantly, while data were collected across three scan sites, exhaustive measures were taken to harmonize acquisitions, including identical scanner hardware, software, and acquisition sequences, as well as comprehensive, standardized staff trainings^26^.

T1- and T2-FLAIR weighted images were obtained from the UK BioBank portal (application 20904) and processed with FreeSurfer 6.0.1. We incorporated all scans that were defined as ‘usable’ by the UK Biobank’s in-house quality control^26^ and completed FreeSurfer processing within 20 hours. ‘Recon-all’ processing included bias field correction, registration to stereotaxic space, intensity normalization, skull-stripping, and segmentation. T2-FLAIR images were used to improve pial surface reconstruction when available. A triangular surface tessellation fitted a deformable mesh model onto the white matter volume, providing grey-white and pial surfaces with >160,000 corresponding vertices registered to fsaverage standard space. Surface area, thickness, and volumetric features were obtained for each of 68 cortical regions (34 per hemisphere) in the Desikan-Killiany atlas^45^ from the aparc.stats files output by the ‘recon-all’ pipeline. Global cerebrum tissue volumes were extracted from the aseg.stats files output by the recon-all process: ‘Total cortical gray matter volume’ for GMV; ‘Total cerebral white matter volume’ for WMV; and ‘Subcortical gray matter volume’ for sGMV (inclusive of thalamus, caudate nucleus, putamen, pallidum, hippocampus, amygdala, and nucleus accumbens area; https://freesurfer.net/fswiki/SubcorticalSegmentation).

Analyses were restricted to individuals who had never had mental health problems as diagnosed by a mental health professional, per their response recorded in data-field 20544 (https://biobank.ndph.ox.ac.uk/showcase/field.cgi?id=20544) of the UKB mental health questionnaire. We further restricted the age range of our sample to subjects ages 50 to 80 years, removing 411 individuals. This was done to reduce the possibility that inadequate representation of both males and females at each end of the sample would induce age differences between the synthetic sites.

#### Software

Subject sampling, main analyses, and visualizations were done in R v 4.1.1. ComBat harmonization, sampling, and analyses for replications were done in R v 4.1.2.

#### Synthetic Site Assignments

##### Main analyses

For our main analyses, each UKB subject in our sample was assigned to one of three synthetic sites using random sampling without replacement in which the probability of a subject being assigned to a given site depended on sex. Female subjects were assigned a 33%, 58.75%, and 8.25% chance of being assigned to Sites A, B, and C, respectively; male subjects had a 33%, 8.25%, and 58.75% likelihood of assignment across these same three sites. Thus the sampling was constructed to create artificial sites of roughly equivalent size but with Male:Female ratios of 1:1 in Site A, 1:4 in Site B, and 4:1 in Site C. This sampling was permuted 100 times for our replication analyses.

##### Varying Male:Female Ratios

In addition, we assessed how the relative performance of ComBat methods varied with varying degrees of imbalance in sites’ Male:Female ratio by reassigning subjects to one of two synthetic sites: one with an equal number of males and females, and one with a sex ratio that varied across 11 permutations. Each site was limited to N=9,400 subjects, so that the 14,213 female UKB participants, who were a slight minority in the sample, could be assigned to one sex-balanced site and one entirely female site without replacement. We used R’s slice_sample() function to assign the appropriate number of randomly selected males and females to each site, with the number of males assigned to the second site increasing by 940 (10% of the site’s sample) from zero to 9,400 over 11 permutations.

#### ComBat Harmonization

For each sampling permutation, we harmonized across the synthetic sites, or batches, using 4 ComBat methods: ComBat, ComBat-GAM, ComBatLS, and linear ComBat without any covariate preservation. Harmonization was performed in R using the ComBatFamily package (https://github.com/andy1764/ComBatFamily). ComBat was fit while accounting for linear effects of age and sex while ComBat-GAM was fit with a thin plate regression spline for age and linear effect of sex. For ComBatLS, the first moment was fit with a penalized b-spline for age and linear effect of sex and the second moment was fit with linear effects for both sex and age. Empirical Bayesian estimation was used within each category of features – global tissue volumes and regional cortical volumes, thicknesses, and surface areas – to improve site effect estimates.

#### Centile score calculation

For our simulations in UKB, we fit simple brain chart models of each feature using GAMLSS such that:

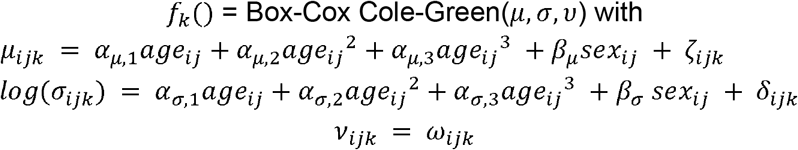

As above, age is calculated in days, and sex is binarized with the reference level as ‘female’. We chose the default Box-Cox Cole-Green distribution method to allow for z-score estimation (see Supplement Section IV).

This model was fit on every feature’s true, unharmonized distribution and on data harmonized with each ComBat method using the *gamlss* package. For each model, we obtained subjects’ centile scores using the *predictAll()* function. This resulted in centile scores for each subject on all features’ true values, as well as their values when harmonized by the four ComBat methods.

#### Evaluating relative accuracy across ComBat methods

To assess the ability of each ComBat method to preserve biologically relevant sources of interindividual variability, even when they are distributed unequally across sites, we tested how closely subjects’ centile scores matched “ground-truth” centiles derived from unharmonized data. This within-subject accuracy was quantified by subtracting the ground-truth centile from the centile derived from harmonized data for each feature, referred to as “centile error”. Negative centile scores indicate that the ComBat method caused an underestimation of the centile score, while positive scores indicate ComBat-induced centile inflation.

As our primary goal was to assess the overall efficacy of each harmonization method, we evaluated the magnitude of inaccuracy within each feature using pairwise comparisons of the absolute values of centile errors between methods. As these distributions were paired, skewed, and unequal in variance, we converted the absolute centile errors into ranks before conducting paired, two-tailed *t*-tests with Welch’s correction^46,47^. Significantly smaller absolute centile errors indicated that a ComBat method preserved covariate effects in that feature more accurately than the ComBat method with larger absolute centile errors. Within each sampling replication, we applied FDR correction across 6 pairings of the four ComBat methods and 208 features for a total of 1,248 comparisons. We applied these same methods to evaluate ComBat methods’ accuracy in our 11 samples with varying male:female ratios, using FDR correction to account for 1,248 comparisons * 11 male:female ratios simulated.

#### Assessing sex-biases in centile scores induced by ComBat methods

We first visualized the effect of sex for each feature’s scale. We used the drop1() function to assess the significance of the sex term in sigma for each feature’s brain charts, as recommended^48^. We then defined the effect of sex on variance as the difference in variance predicted for a male minus the variance predicted for a female at the average age of the sample, 64.94 years. We calculated a standardized effect of sex that would be comparable across brain features by dividing this sex effect by the predicted variance for females (the reference level for sex).

To determine whether ComBat harmonization differentially affected males and females when applied to sites with imbalanced sex samples, we conducted within-feature, two-tailed *t*-tests of ranks with Welch’s correction^46,47^ comparing males’ and female’s centile errors. These analyses were corrected for comparisons of four ComBat methods across 208 features using FDR adjustment and were repeated across our 100 sampling replications as well as 11 simulated male:female ratios. We defined the size of this sex-effect within a feature as the male centile error’s median minus the female centile error’s median, which produces a single sex-difference metric.

We conducted several further analyses to determine whether sex-differences were systematic across features such that centile scores would be biased in a particular direction.

First, summarized sex differences within each feature type – global tissue volumes, regional cortical volumes, regional cortical thickness, and regional surface area – by taking the median of their sex-difference metrics. Finally, we repeated this procedure across our 100 sampling replications to create a distribution of median sex differences for each ComBat method and feature type, then used one-sample *t*-tests to assess whether these distributions’ means differed from zero. A non-zero mean would indicate that, for a given feature type, that ComBat method tends to bias centile scores by sex, with more negative distributions indicating that males’ centiles tend to be underestimated relative to females’. Analyses were corrected for 16 comparisons using FDR adjustment.

Next, we determined whether ComBat-induced errors led to one sex being overrepresented among those with extreme centile scores, which could mask or skew associations between centiles and other phenotypes of interest. We first calculated subjects’ average centile scores within global tissue volume, regional cortical volume, regional cortical thickness, and regional surface area features. We considered average centiles below 20% as “low” and above 80% as “high”. We used centiles derived from the true, unharmonized data to calculate what proportion of individuals in each category was expected to be female. Finally, we compared this expected proportion to the distribution of proportions calculated across 100 replications of harmonized data using two-sided, one-sample *t*-tests, FDR corrected across four ComBat methods, four phenotype categories, and two high/low groupings.

Finally, we assessed directional sex-biases across our 11 sampling permutations with varying sex ratios by calculating the sex difference in medians for each feature’s centile errors, as in our main analyses. We first used within-feature, two-tailed *t*-tests of ranks with Welch’s correction to compare males’ and females’ centile errors using FDR correction, as above. We then used one-sample *t*-tests to assess whether the mean of these significant within-feature sex differences differed from zero in any of the sex-imbalanced permutations tested. FDR was corrected for testing sex-difference distributions from 4 ComBat methods across 11 levels of sex-imbalanced sampling.

### Harmonization of neuroanatomical features from the LBCC dataset

#### Lifespan Brain Chart Consortium sample

The Lifespan Brain Chart Consortium (LBCC) is an aggregated collection of structural MRI scans including data from individuals across the range of the human lifespan and the globe. Details of the dataset and primary studies, including diagnostic criteria, can be found elsewhere^20^. For these analyses, we used data from healthy individuals collected by 60 studies. We first excluded all subjects under age of 3, removing a total of 728 individuals. We then assessed image quality using the Euler index, an automated measure of reconstructions’ surface continuity often used as a robust, quantitative assessment of scan quality^49^. As in prior work^20^, we applied an adaptive threshold of the Euler index, excluding scans with Euler indices greater than two median absolute deviations above the primary-study median. This quality threshold removed a total of 6,282 scans (8.46% of a study’s primary sample on average), while a further 1981 subjects with missing values were removed via listwise deletion. Following quality control procedures, a total of 52,098 scans from 51 studies and 199 sites remained for analysis.

#### ComBat Harmonization

We applied ComBat-GAM and ComBatLS to harmonize this curated LBCC sample, treating primary collection sites as batches. Harmonization was performed on log-transformed feature values to prevent ComBat from estimating negative feature values. We preserved the effects of age, sex, and their interaction by estimating as follows study **i**, subject **j**, and feature **k**:

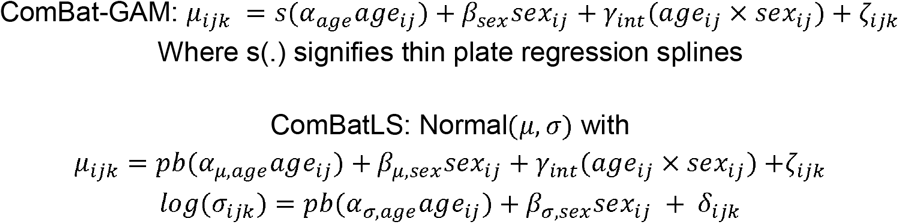

Where pb(.) signifies penalized b-splines, with 20 knots allowed in the mu term and 5 knots in sigma. Models that failed to converge using the *gamlss* package-default RS() algorithm were attempted using CG()^48^.

Empirical Bayesian estimation was used within each category of cortical features – volumes, thickness, and surface area – to improve site effect estimates, while effects for each global tissue volume were estimated individually.

#### Centile score calculation

We used ComBatLS and ComBat-GAM harmonized datasets as well as raw data to estimate each feature’s normative trajectory across the LBCC sample with the following GAMLSS model:

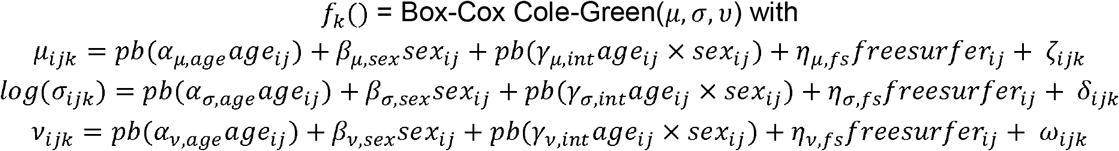

where pb(.) signifies penalized b-splines. As above, age is calculated in days, and sex is binarized with the reference level as ‘female’. We also controlled for variation in image-processing software (‘freesurfer’) and the corresponding sums of all brain volume, cortical gray matter volume, cortical thickness, and surface area features when modeling global tissue volumes and regional cortical measures (‘global’).

Models failed to converge for two features in the ComBatLS dataset, and one feature in the ComBat-GAM dataset, and one feature in the raw dataset, for a total of two features excluded from further analysis. Subjects’ centile scores were derived from each model using the predictAll() function.

#### Site-Effect Quantification

We conducted two tests to assess how well ComBatLS harmonizes real world data, using data harmonized by ComBat-GAM as a “silver standard” and the raw data as a lower bound. Within each dataset, calculated each subjects’ average centile score across all available features, then used ANOVA to assess whether a subjects’ average centiles varied across primary studies. To directly quantify the residual effects of primary study within each feature, we re-fit the gamlss brain chart models specified above on the ComBatLS- and ComBat-GAM-harmonized datasets, this time including fixed effects of study in the mu and sigma moments. From there, we used generalized pseudo R-squared^48,50^ to calculate the local effect size of “study” via Cohen’s F-squared^51^.

#### Comparison of ComBatLS and ComBat-GAM and associations with sites’ sample characteristics

While no “ground truth” for harmonized LBCC data exists, we evaluated whether differences in subjects’ centile scores when harmonized with ComBatLS and ComBat-GAM were related to the distribution of biological sources of variability across studies. Specifically, as studies tended only contain subjects from a relatively narrow age range, we assessed whether a subject’s average magnitude of difference in ComBatLS- and ComBat-GAM-derived centiles was associated with the mean age of its primary study or how greatly that individual deviated from their primary study’s mean age. For each subject, we calculated differences in centiles from the two methods and took the mean of their absolute values across features. We also calculated the mean age of each site’s sample and the absolute difference between that mean age and ages of each individual in the study sample. Finally, we used linear regression to test subjects’ mean absolute centile differences against the study’s mean age and their absolute deviation from that mean age, respectively, while controlling for study sample size.

## Acknowledgments

MG, LD, JS, and AAB are supported by R01MH134896, R01MH132934, and R01MH133843 from the National Institutes of Health. MG, RTS, SS, JS, and AAB are supported by “RAising the Investment in Sex and gender Evidence (RAISE) Pilot Grant Award” funded by FOCUS on Health and Leadership for Women and Penn PROMOTES Research on Sex and Gender in Health. SS was supported by DP5OD036142 from the NIH Common Fund, a 2022 Young Investigator Grant from the Brain & Behavior Research Foundation, and a 2023 Career Award for Medical Scientists from the Burroughs Wellcome Fund. RTS is funded by R01MH123550, R01MH112847, R01MH123563, R01NS112274, and R01NS060910. RRG is funded by the EMERGIA Junta de Andalucía program (EMERGIA20_00139), the Plan de Consolidación (CNS2023-143647) and the Plan de Generación de Conocimiento from the Agencia Estatal de Investigación (PID2021-122853OA-I00).

## References

1. Marek, S. et al. Reproducible brain-wide association studies require thousands of individuals. Nature 603, 654 (2022).

2. Smith, S. M. & Nichols, T. E. Statistical Challenges in “Big Data” Human Neuroimaging. Neuron 97, 263–268 (2018).

3. Thompson, P. M. et al. ENIGMA and global neuroscience: A decade of large-scale studies of the brain in health and disease across more than 40 countries. Transl. Psychiatry 10, 1–28 (2020).

4. Bayer, J. M. M. et al. Site effects how-to and when: An overview of retrospective techniques to accommodate site effects in multi-site neuroimaging analyses. Front. Neurol. 13, (2022).

5. Hu, F. et al. Image harmonization: A review of statistical and deep learning methods for removing batch effects and evaluation metrics for effective harmonization. NeuroImage 274, 120125 (2023).

6. Johnson, W. E., Li, C. & Rabinovic, A. Adjusting batch effects in microarray expression data using empirical Bayes methods. Biostat. Oxf. Engl. 8, 118–127 (2007).

7. Fortin, J. P. et al. Harmonization of cortical thickness measurements across scanners and sites. NeuroImage 167, 104 (2018).

8. Fortin, J.-P. et al. Harmonization of multi-site diffusion tensor imaging data. NeuroImage 161, 149–170 (2017).

9. Pomponio, R. et al. Harmonization of large MRI datasets for the analysis of brain imaging patterns throughout the lifespan. NeuroImage 208, (2020).

10. Orlhac, F. et al. A Guide to ComBat Harmonization of Imaging Biomarkers in Multicenter Studies. J. Nucl. Med. 63, 172–179 (2022).

11. Yu, M. et al. Statistical harmonization corrects site effects in functional connectivity measurements from multi-site fMRI data. Hum. Brain Mapp. 39, 4213–4227 (2018).

12. Sun, D. et al. A comparison of methods to harmonize cortical thickness measurements across scanners and sites. NeuroImage 261, 119509 (2022).

13. Radua, J. et al. Increased power by harmonizing structural MRI site differences with the ComBat batch adjustment method in ENIGMA. NeuroImage 218, 116956 (2020).

14. Forde, N. J. et al. Sex Differences in Variability of Brain Structure Across the Lifespan. Cereb. Cortex N. Y. NY 30, 5420 (2020).

15. Wierenga, L. M. et al. Greater male than female variability in regional brain structure across the lifespan. Hum. Brain Mapp. 43, 470 (2022).

16. Wierenga, L. M., Sexton, J. A., Laake, P., Giedd, J. N. & Tamnes, C. K. A Key Characteristic of Sex Differences in the Developing Brain: Greater Variability in Brain Structure of Boys than Girls. Cereb. Cortex N. Y. NY 28, 2741–2751 (2018).

17. Dickie, D. A. et al. Variance in Brain Volume with Advancing Age: Implications for Defining the Limits of Normality. PLoS ONE 8, e84093 (2013).

18. Lange, N., Giedd, J. N., Castellanos, F. X., Vaituzis, A. C. & Rapoport, J. L. Variability of human brain structure size: ages 4-20 years. Psychiatry Res. 74, 1–12 (1997).

19. Williams, C. M., Peyre, H., Toro, R. & Ramus, F. Neuroanatomical norms in the UK Biobank: The impact of allometric scaling, sex, and age. Hum. Brain Mapp. 42, 4623–4642 (2021).

20. Bethlehem, R. A. I. et al. Brain charts for the human lifespan. Nat. 2022 6047906 604, 525–533 (2022).

21. Dima, D. et al. Subcortical volumes across the lifespan: Data from 18,605 healthy individuals aged 3–90 years. Hum. Brain Mapp. 43, 452–469 (2021).

22. Frangou, S. et al. Cortical thickness across the lifespan: Data from 17,075 healthy individuals aged 3-90 years. Hum. Brain Mapp. 43, 431–451 (2022).

23. Rutherford, S. et al. Evidence for embracing normative modeling. eLife 12, e85082 (2023).

24. Nygaard, V., Rødland, E. A. & Hovig, E. Methods that remove batch effects while retaining group differences may lead to exaggerated confidence in downstream analyses. Biostatistics 17, 29–39 (2016).

25. Rigby, R. A. & Stasinopoulos, D. M. Generalized additive models for location, scale and shape. J. R. Stat. Soc. Ser. C Appl. Stat. 54, 507–554 (2005).

26. Littlejohns, T. J. et al. The UK Biobank imaging enhancement of 100,000 participants: rationale, data collection, management and future directions. Nat. Commun. 11, 2624 (2020).

27. Bach, S., Morrow, M. M., Zhao, K. D. & Hughes, R. E. Sex Distribution of Study Samples Reported in American Society of Biomechanics Annual Meeting Abstracts. PLoS ONE 10, e0118797 (2015).

28. Rechlin, R. K., Splinter, T. F. L., Hodges, T. E., Albert, A. Y. & Galea, L. A. M. An analysis of neuroscience and psychiatry papers published from 2009 and 2019 outlines opportunities for increasing discovery of sex differences. Nat. Commun. 13, 2137 (2022).

29. Dickinson, E. R., Adelson, J. L. & Owen, J. Gender Balance, Representativeness, and Statistical Power in Sexuality Research Using Undergraduate Student Samples. Arch. Sex. Behav. 41, 325–327 (2012).

30. Ritchie, S. J. et al. Sex Differences in the Adult Human Brain: Evidence from 5216 UK Biobank Participants. Cereb. Cortex 28, 2959–2975 (2018).

31. Marquand, A. F. et al. Conceptualizing mental disorders as deviations from normative functioning. Mol. Psychiatry 24, 1415–1424 (2019).

32. Lombardi, A. et al. Extensive Evaluation of Morphological Statistical Harmonization for Brain Age Prediction. Brain Sci. 10, 364 (2020).

33. Yu, Y. et al. Brain-Age Prediction: Systematic Evaluation of Site Effects, and Sample Age Range and Size. 2023.11.06.565917 Preprint at 10.1101/2023.11.06.565917 (2023).

34. Kang, K. et al. Study design features that improve effect sizes in cross-sectional and longitudinal brain-wide association studies. bioRxiv doi:10.1101/2023.05.29.542742.

35. Zhang, Y., Jenkins, D. F., Manimaran, S. & Johnson, W. E. Alternative empirical Bayes models for adjusting for batch effects in genomic studies. BMC Bioinformatics 19, 262 (2018).

36. Gebre, R. K. et al. Cross–scanner harmonization methods for structural MRI may need further work: A comparison study. NeuroImage 269, 119912 (2023).

37. Bayer, J. M. M. et al. Accommodating site variation in neuroimaging data using normative and hierarchical Bayesian models. NeuroImage 264, 119699 (2022).

38. Bridgeford, E. et al. A Causal Perspective for Batch Effects: When Is No Answer Better than a Wrong Answer? (2024). doi:10.1101/2021.09.03.458920.

39. Zhu, A. H. et al. Reference curves for harmonizing multi-site regional diffusion MRI metrics across the lifespan. Preprint at 10.1101/2024.02.22.581646v1 (2024).

40. de Boer, A. A. A. et al. Non-Gaussian normative modelling with hierarchical Bayesian regression. Imaging Neurosci. 2, 1–36 (2024).

41. Beery, A. K. Inclusion of females does not increase variability in rodent research studies. Curr. Opin. Behav. Sci. 23, 143–149 (2018).

42. Zajitschek, S. R. et al. Sexual dimorphism in trait variability and its eco-evolutionary and statistical implications. eLife 9, e63170.

43. Khodursky, S. et al. Sex differences in interindividual gene expression variability across human tissues. PNAS Nexus 1, pgac243 (2022).

44. Harvey, A. C. Estimating Regression Models with Multiplicative Heteroscedasticity. Econometrica 44, 461–465 (1976).

45. Desikan, R. S. et al. An automated labeling system for subdividing the human cerebral cortex on MRI scans into gyral based regions of interest. NeuroImage 31, 968–980 (2006).

46. Zimmerman, D. W. & Zumbo, B. D. Rank transformations and the power of the Student t test and Welch t’ test for non-normal populations with unequal variances. Can. J. Exp. Psychol. Rev. Can. Psychol. Expérimentale 47, 523–539 (1993).

47. Ruxton, G. D. The unequal variance t-test is an underused alternative to Student’s t-test and the Mann–Whitney U test. Behav. Ecol. 17, 688–690 (2006).

48. Stasinopoulos, D. M. & Rigby, R. A. Generalized Additive Models for Location Scale and Shape (GAMLSS) in R. J. Stat. Softw. 23, 1–46 (2008).

49. Rosen, A. F. G. et al. Quantitative assessment of structural image quality. NeuroImage 169, 407–418 (2018).

50. Nagelkerke, N. J. D. A note on a general definition of the coefficient of determination. Biometrika 78, 691–692 (1991).

51. Selya, A. S., Rose, J. S., Dierker, L. C., Hedeker, D. & Mermelstein, R. J. A Practical Guide to Calculating Cohen’s f2, a Measure of Local Effect Size, from PROC MIXED. Front. Psychol. 3, (2012).

