## Supplementary material for "ComBatLS: A location- and scale-preserving method for multi-site image harmonization": combatls_supplement_preprint.pdf

**Contents**


I. **Results including ComBat without covariate preservation**
II. **The magnitude of sex's effect on variance depends on the type of brain feature measured**
III. **Replication without Extreme Centiles**
IV. **Z-score analyses**
   A. **Main Analyses**
   B. **Varying M:F ratios**
   C. **Replication without Extreme Z-scores**
V. **Comparisons of ComBatLS and ComBat-GAM for harmonizing consortium data**


### Section I. Results across 100 sampling replications including ComBat without Covariate Preservation

We included an application of ComBat in which no covariate effects are preserved to serve as a benchmark harmonization method against which we could assess the affects of increasingly complex covariate preservation: linear ComBat, ComBat-GAM, and ComBatLS. While tests with this method are controlled for throughout the analyses as part of our FDR corrections, we chose not to include results in the main text to facilitate easy comparison of our primary methods of interest. Results from all four methods are presented below.

Pairwise comparisons of ComBat methods' absolute centile errors for 208 brain features across 100 replications

| Method Producing Smaller Absolute Errors | N Features | % Features |
|---|---|---|
| ComBat-GAM vs ComBat w/o Covariates | | |
| ComBat w/o Covariates | 190 | 0.91% |
| ComBat-GAM | 20605 | 99.06% |
| ComBatLS vs ComBat w/o Covariates | | |
| ComBat w/o Covariates | 155 | 0.75% |
| ComBatLS | 20638 | 99.22% |
| ComBatLS vs ComBat-GAM | | |
| ComBat-GAM | 1503 | 7.23% |
| ComBatLS | 19115 | 91.90% |
| Linear ComBat vs ComBat w/o Covariates | | |
| ComBat w/o Covariates | 190 | 0.91% |
| Linear ComBat | 20604 | 99.06% |
| Linear ComBat vs ComBat-GAM | | |
| ComBat-GAM | 8093 | 38.91% |
| Linear ComBat | 9918 | 47.68% |
| Linear ComBat vs ComBatLS | | |
| ComBatLS | 19146 | 92.05% |
| Linear ComBat | 1466 | 7.05% |

**Supplemental Table 1**) Pairwise tests of absolute centile errors within each of 208 brain features replicated across 100 subject resamplings. All tests conducted as pairwise, two-tailed t-tests of ranks with Welch's correction. FDR-corrected across 1248 tests (208 features x 6 ComBat method pairings) within each sampling permutation.

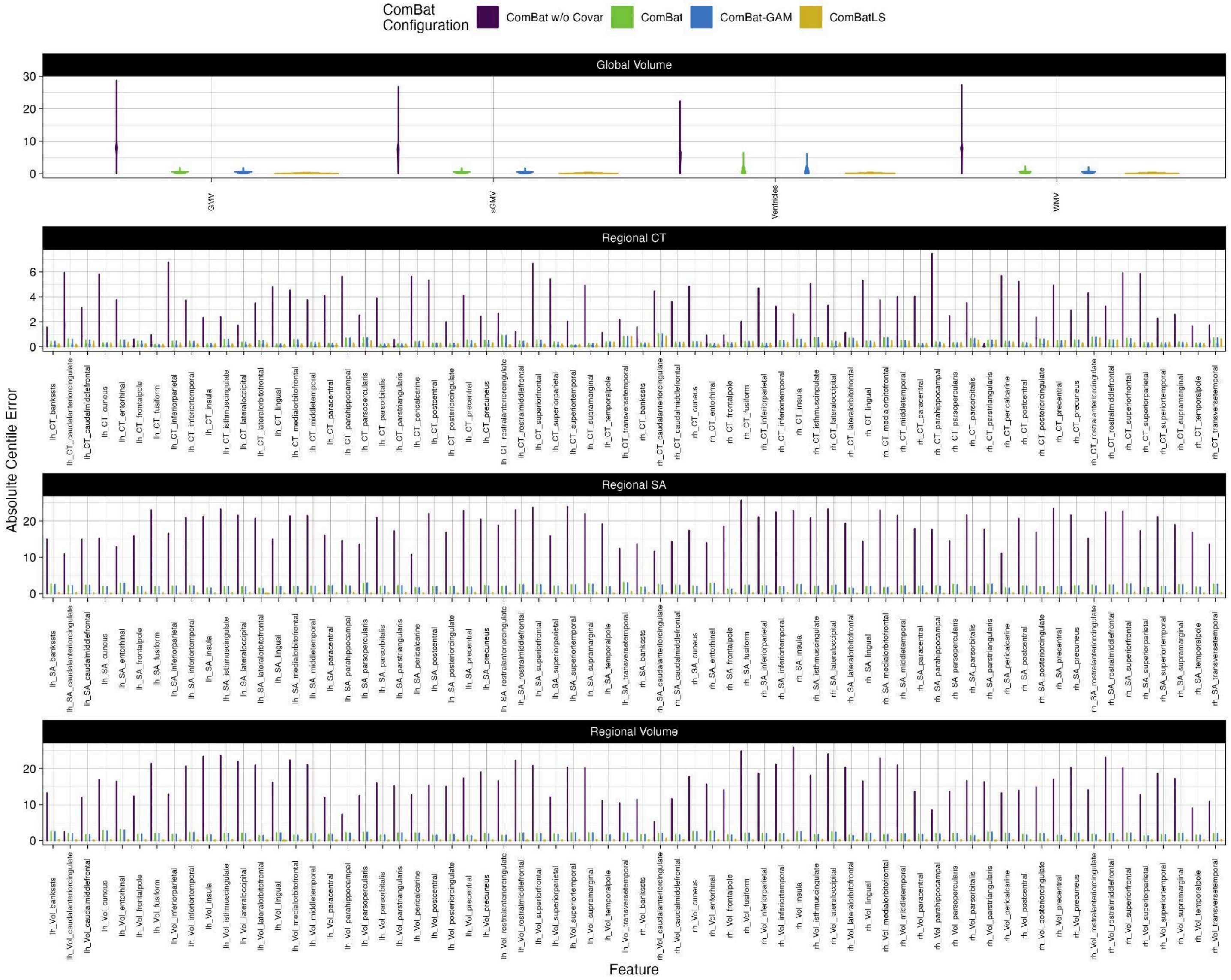


**Supplemental Figure 1. Absolute centile errors across brain features and ComBat methods.** Violin plots of absolute centile errors across 208 brain features.

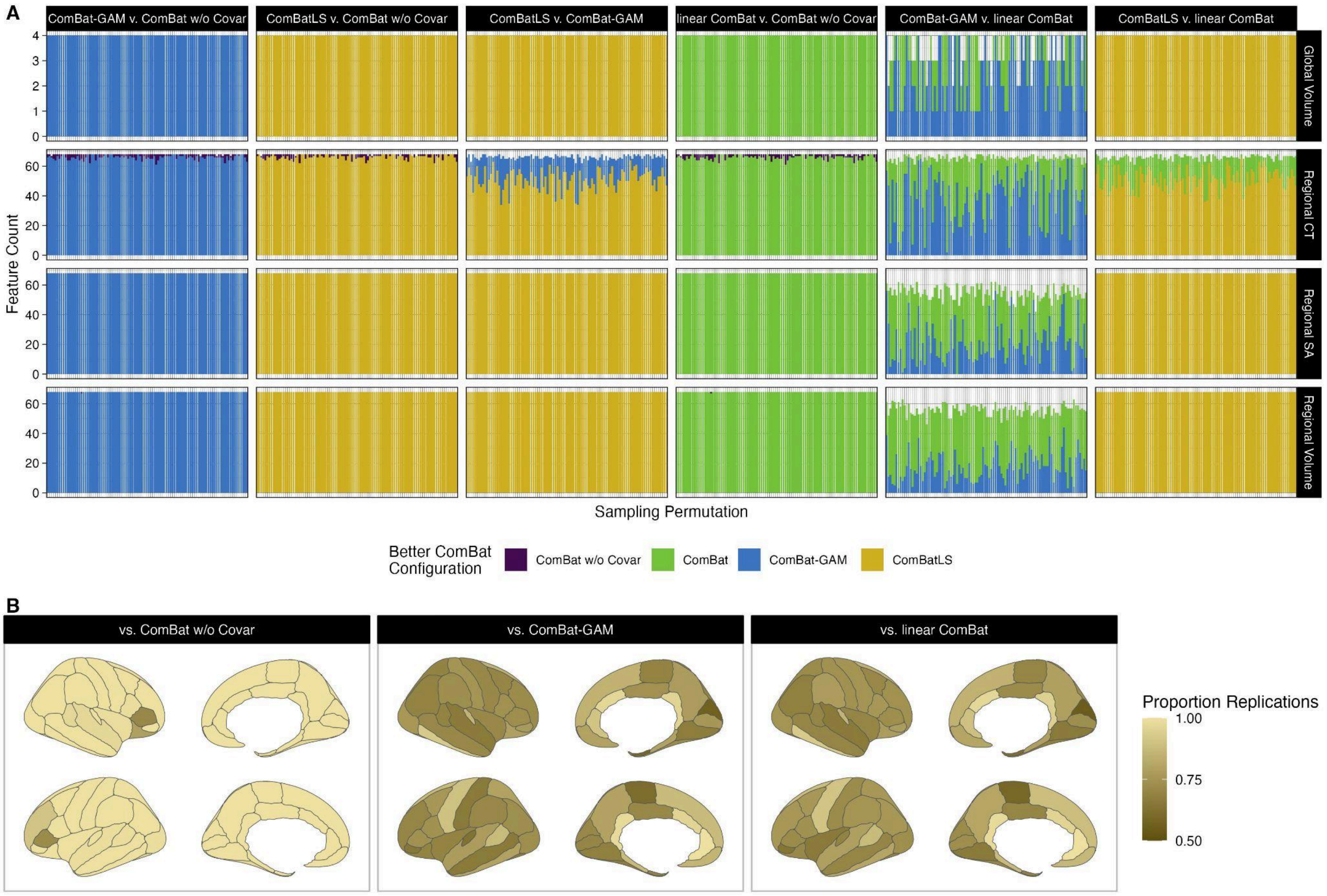


**Supplemental Figure 2. ComBatLS recapitulates true centile scores more accurately than other ComBat methods.** A) Absolute centile errors within each brain feature compared pairwise between four ComBat methods, replicated across 100 sampling permutations. Fill indicates the ComBat method that produces significantly smaller absolute centile errors, FDR-corrected across 1248 tests (208 features x 6 ComBat method pairings) within each permutation. B) Proportion of sampling replications in which ComBatLS produces significantly smaller absolute centile errors for a cortical thickness feature than an alternative ComBat method. Significant differences were assessed using pairwise t-tests between ComBat methods, FDR-corrected. Abbrv: CT, cortical thickness; SA, surface area.

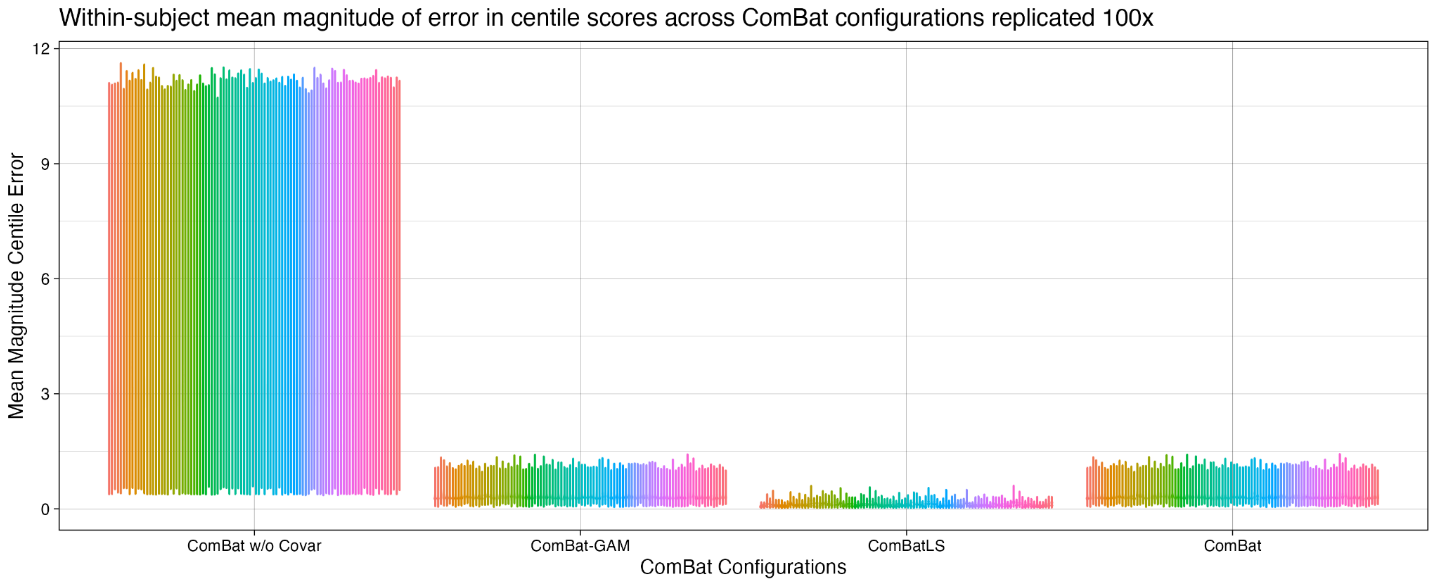


**Supplemental Figure 3. Subjects' mean absolute centile error across ComBat configurations and 100 sampling replications.** Violin plots of absolute centile error for 208 features averaged within subject. Fill corresponds to sampling replication.

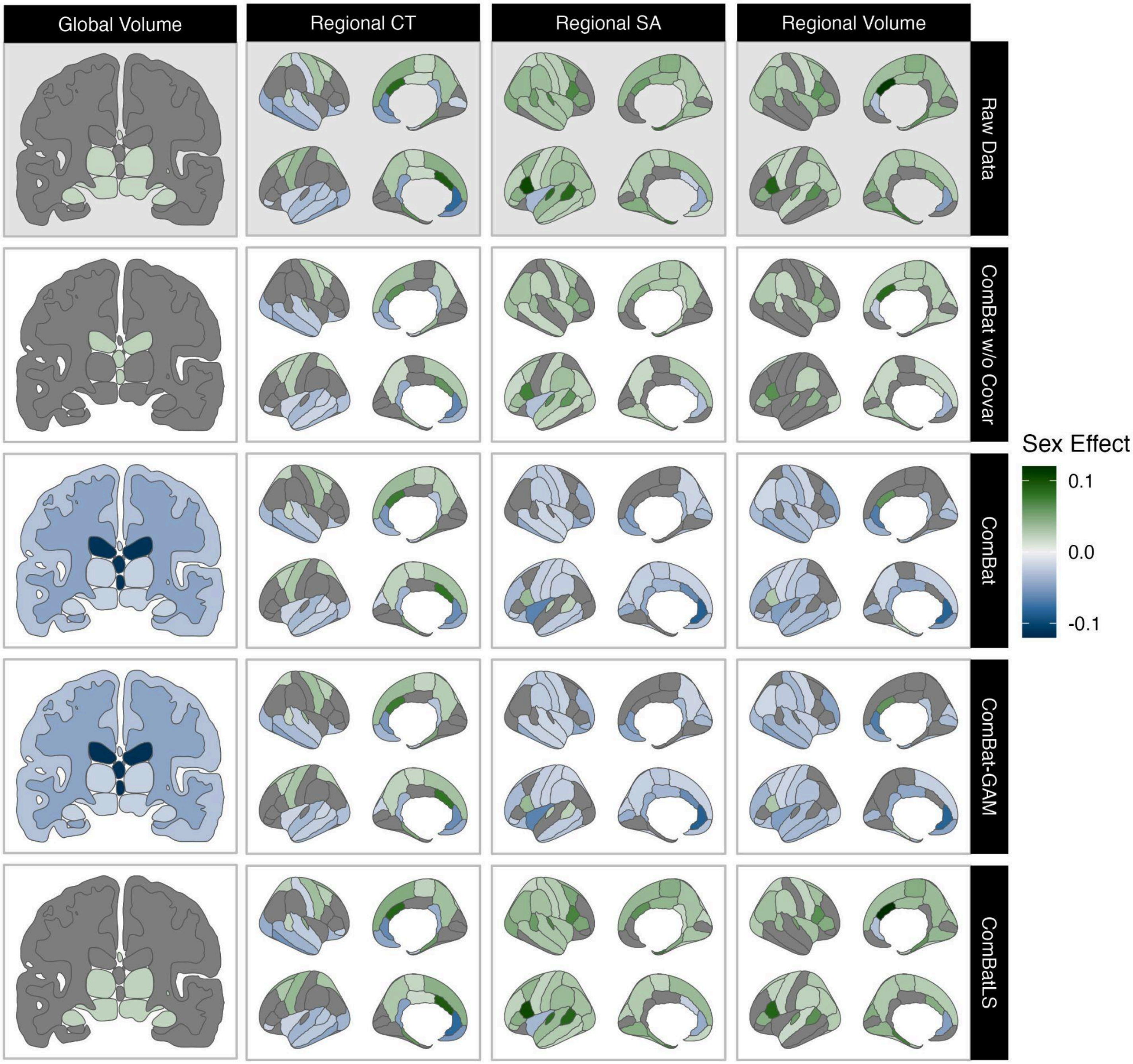


**Supplemental Figure 4. Brain features with significant sex effects in scale**. Features in gray are those for which sex does not significantly impact the second moment of a gamlss brain chart. Fill represents the difference in males' and females' predicted variance at the sample's mean age (64.94 years), standardized by dividing by females' predicted variance. Positive effects indicating that males' variance is higher than females'.

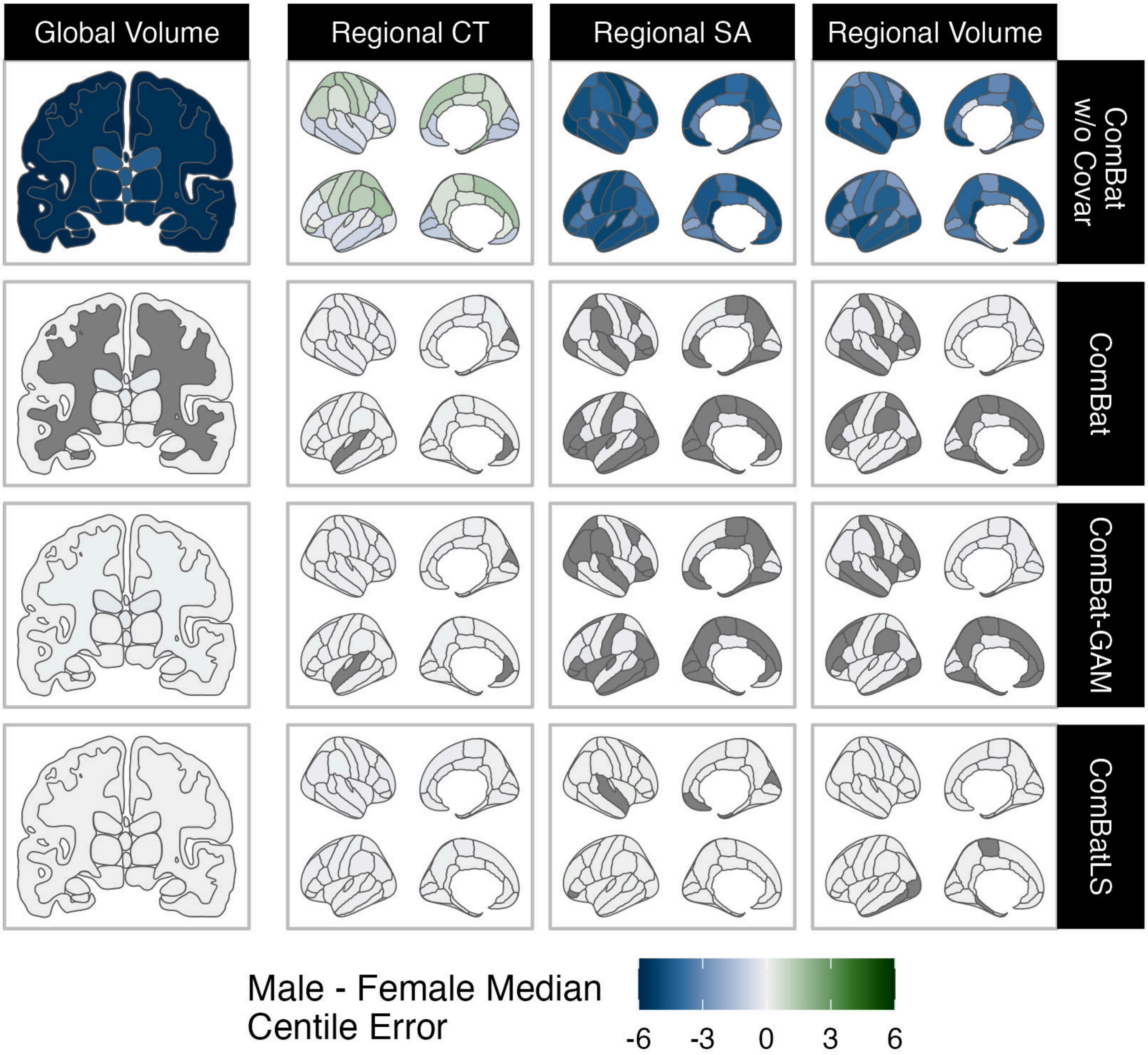


**Supplemental Figure 5. Significant differences in males' and females' median centile errors across brain features and ComBat methods.** Positive centile errors (green) indicate that males' centiles tend to be overestimated relative to females'. Abbrv: CT, cortical thickness; SA, surface area.

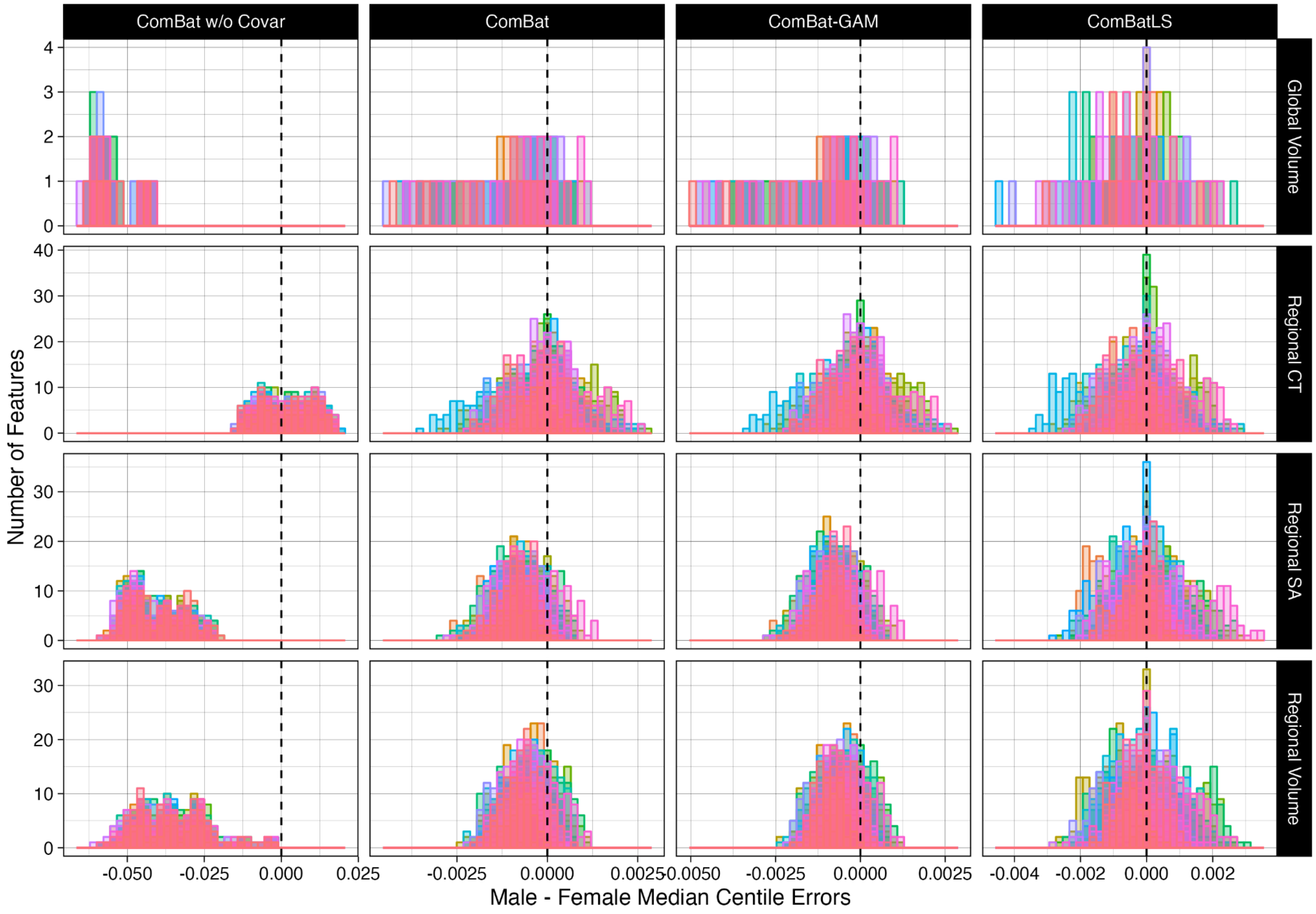


**Supplemental Figure 6. Differences between males' and females' median centile errors in 208 brain features across 100 sampling replications.** Feature counts are plotted by brain tissue types. Fill represents sampling replication. Abbrv: CT, cortical thickness; SA, surface area.

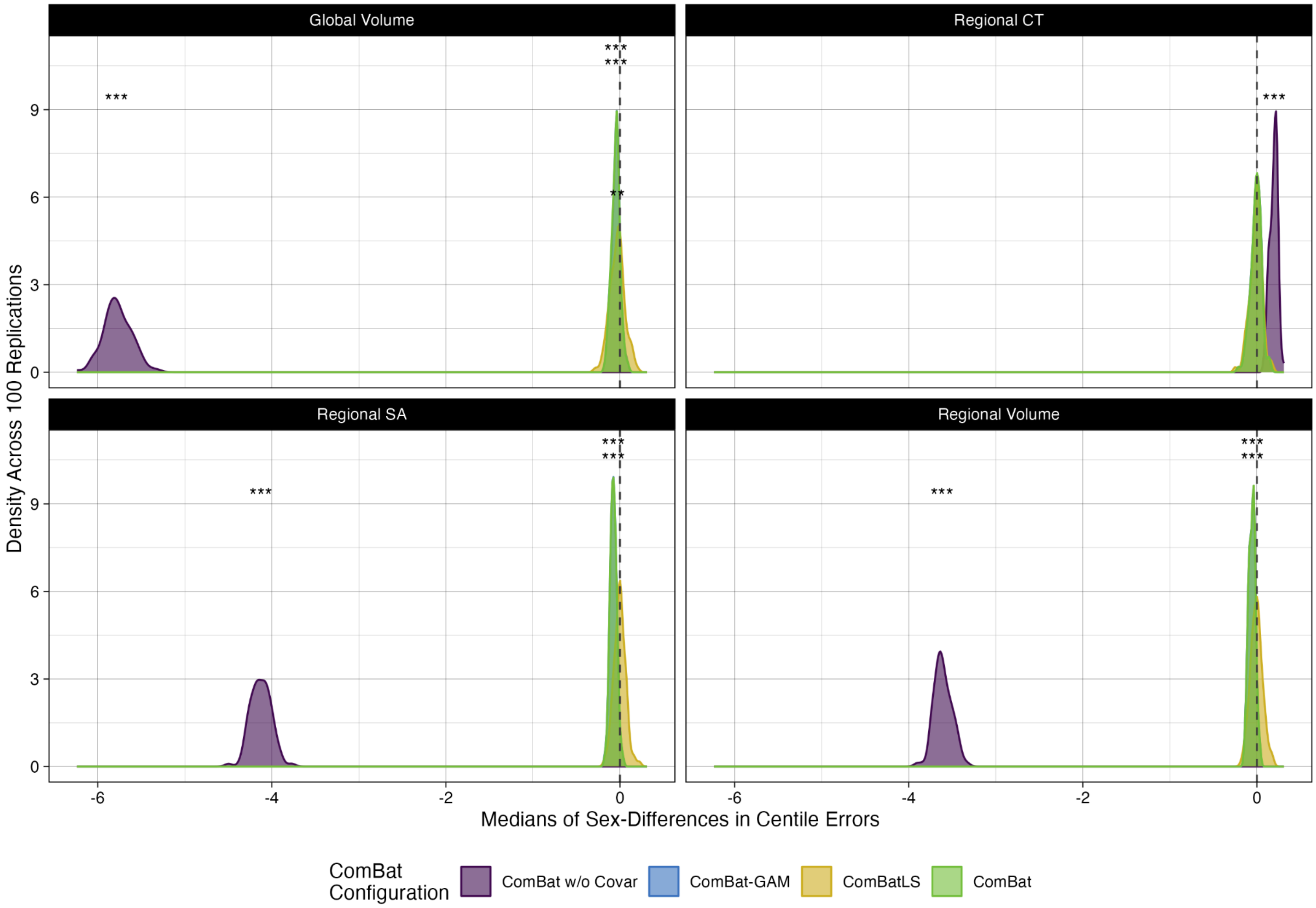


**Supplemental Figure 7. Density plots of median sex differences in centile errors induced by different ComBat methods within phenotype categories across 100 replications.** Data shows medians of the distributions of each replication plotted in Supplemental Figure 5. Abbrv: CT, cortical thickness; SA, surface area; ***, p < 0.001; **, p < 0.01, FDR-corrected.

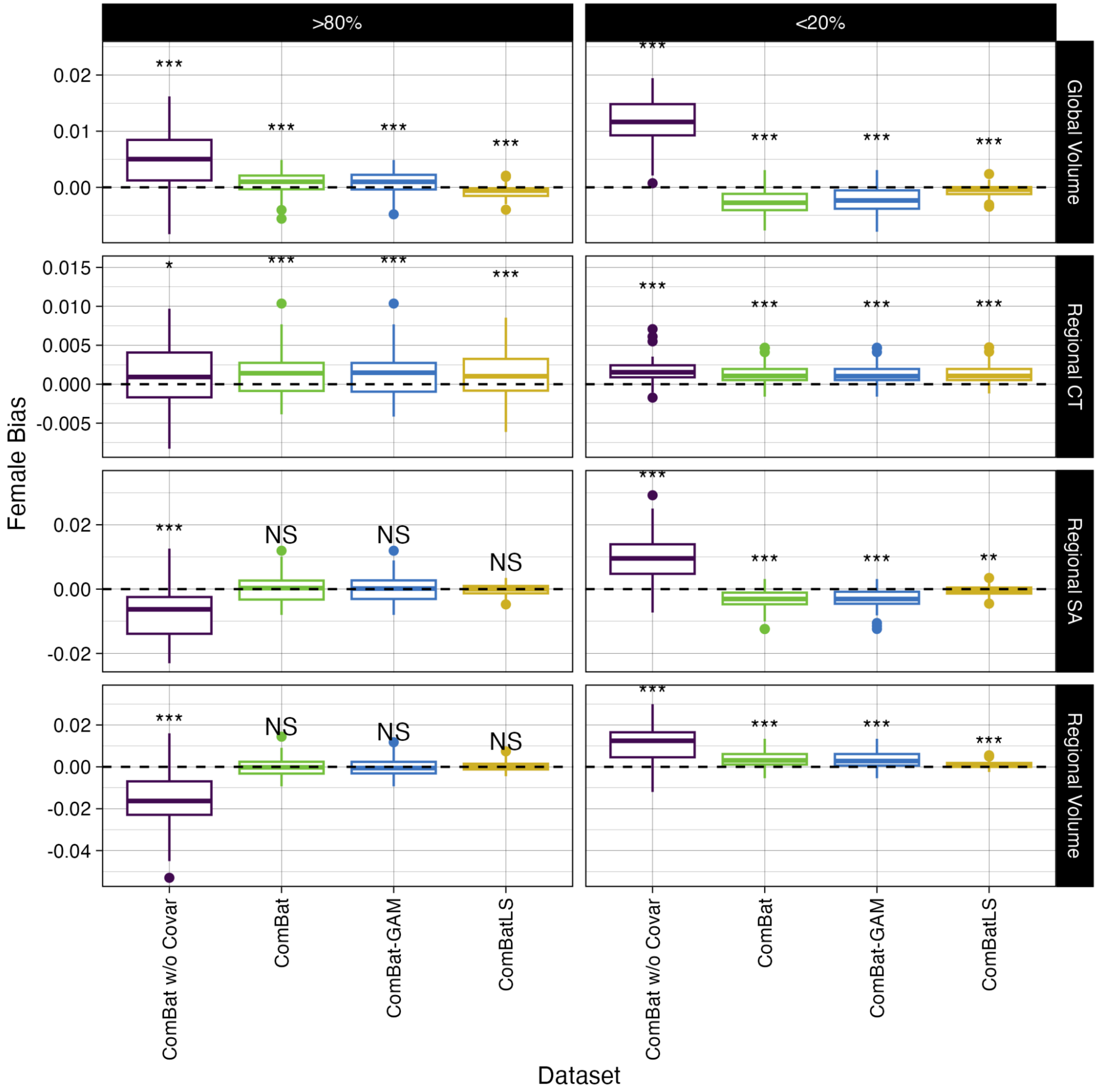


**Supplemental Figure 8. ComBat methods may lead females to be slightly over- or under-represented among individuals with extreme phenotypes.** Bias in the proportion of females with low (<20th percentile) or high (>80th) mean centiles across 100 sampling replications. Positive values indicate a higher proportion of females than “true” mean centiles calculated from unharmonized data (dashed line). Abbrv: ***, $p < 0.001$; **, $p < 0.01$, FDR-corrected.

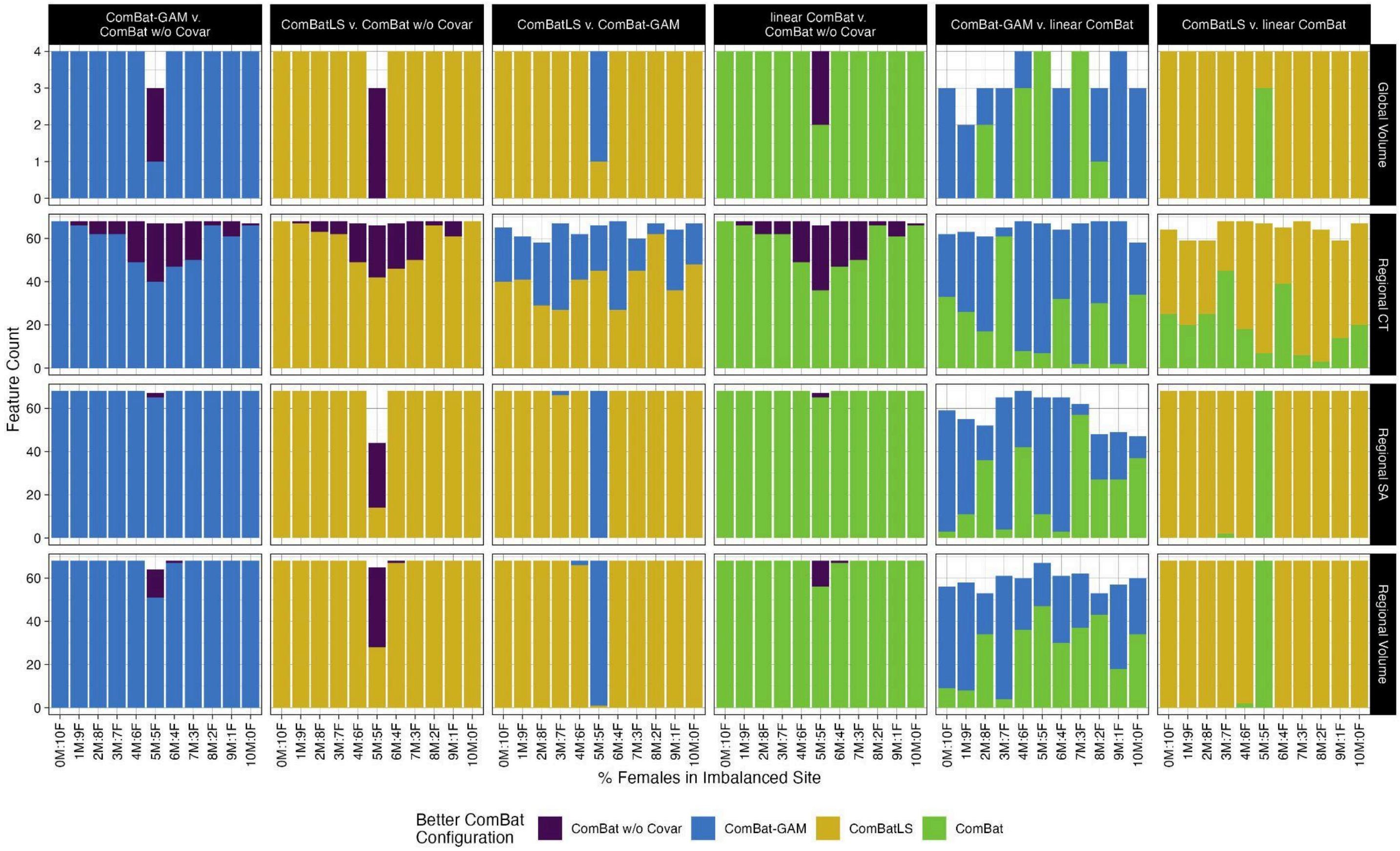


**Supplemental Figure 9. Magnitude centile errors compared pairwise between ComBat methods across varying levels of sex-imbalances in simulated sites.** Fill indicates ComBat method with significantly lower absolute centile errors for a given feature, FDR-corrected across pairwise combinations, brain features, and 11 samplings. Abbrv: CT, cortical thickness; SA, surface area.

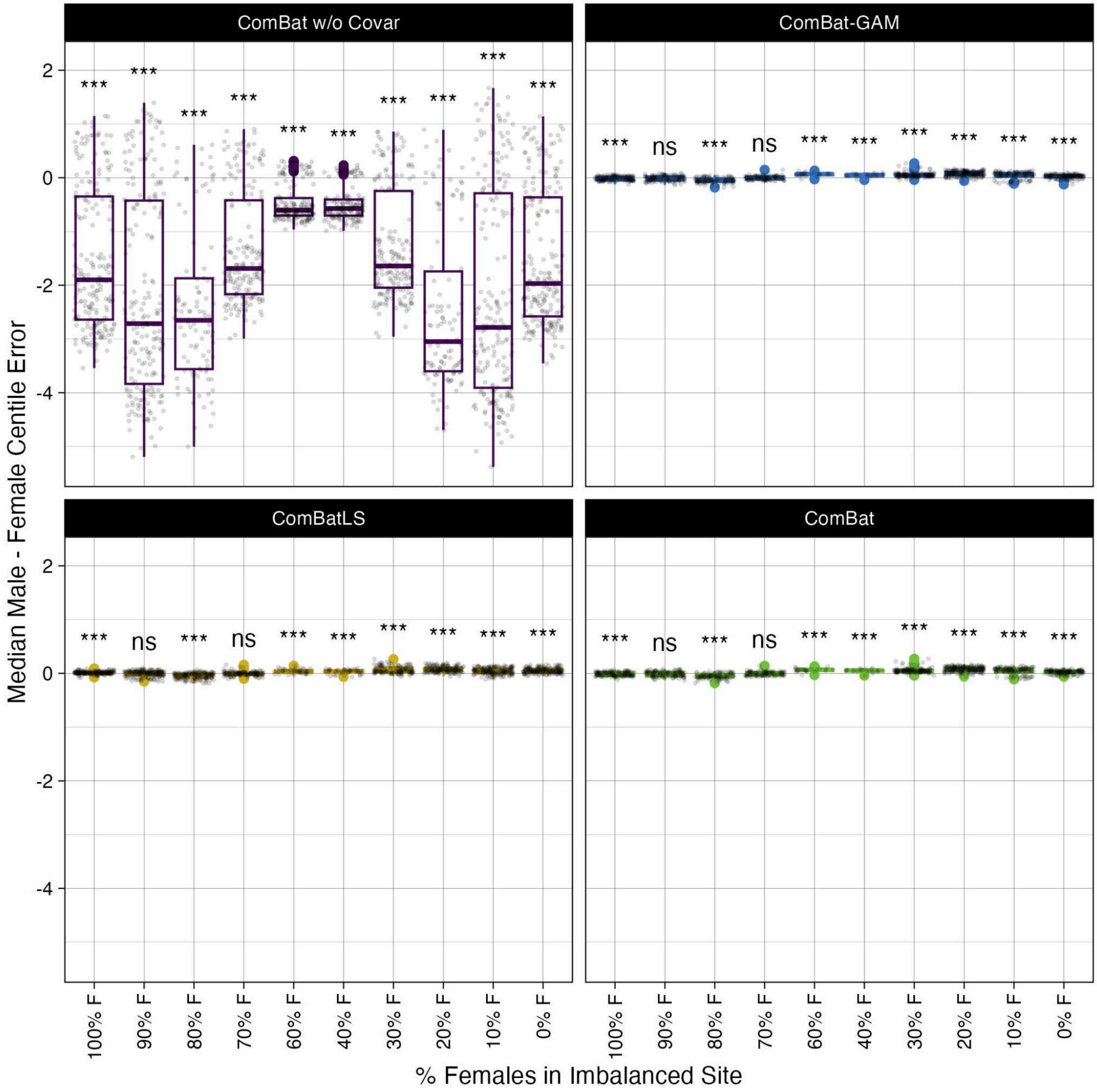


**Supplemental Figure 10. Sex-biases in centile errors induced by various ComBat methods across varying degrees of sex-imbalance.** Points show brain features with significant differences in the distributions of males' and females' centile errors (FDR corrected). Boxplots show median male - median female centile errors across these features when centiles are derived from data harmonized by different ComBat methods. ComBat without covariate preservation induces strong biases wherein males' centiles are underestimated relative to females', particularly as simulated sites become more imbalance for sex. Note that no features display significant sex differences when both sites are balanced for sex (i.e. 50% F). Abbrv: ***, $p < 0.001$; **, $p < 0.01$, FDR-corrected.

**Section II. The magnitude of sex's effect on variance depends on the type of brain feature measured**

Motivated by prior literature, we used UKB data to assess how sex's effects on scale vary by brain feature type. Sex's effects in variance were calculated from brain charts and standardized across features (see Methods). We performed a nonparametric Kruskal-Wallis test to establish that the distibutions of standardized sex effects varied across brain feature types: global tissue volumes (4 features), regional cortical thickness (68 features), regional surface area (68 features), and regional volume (68 features). We then conducted pairwise Wilcox tests (FDR-corrected) which show that cortical thickness features's scales are significantly less impacted by sex than cortical surface area or cortical regional volume features. Sex-effects on cortical thickness features' scales are not significantly smaller than those of global volumes, though this analysis is limited by small number of global volume features.

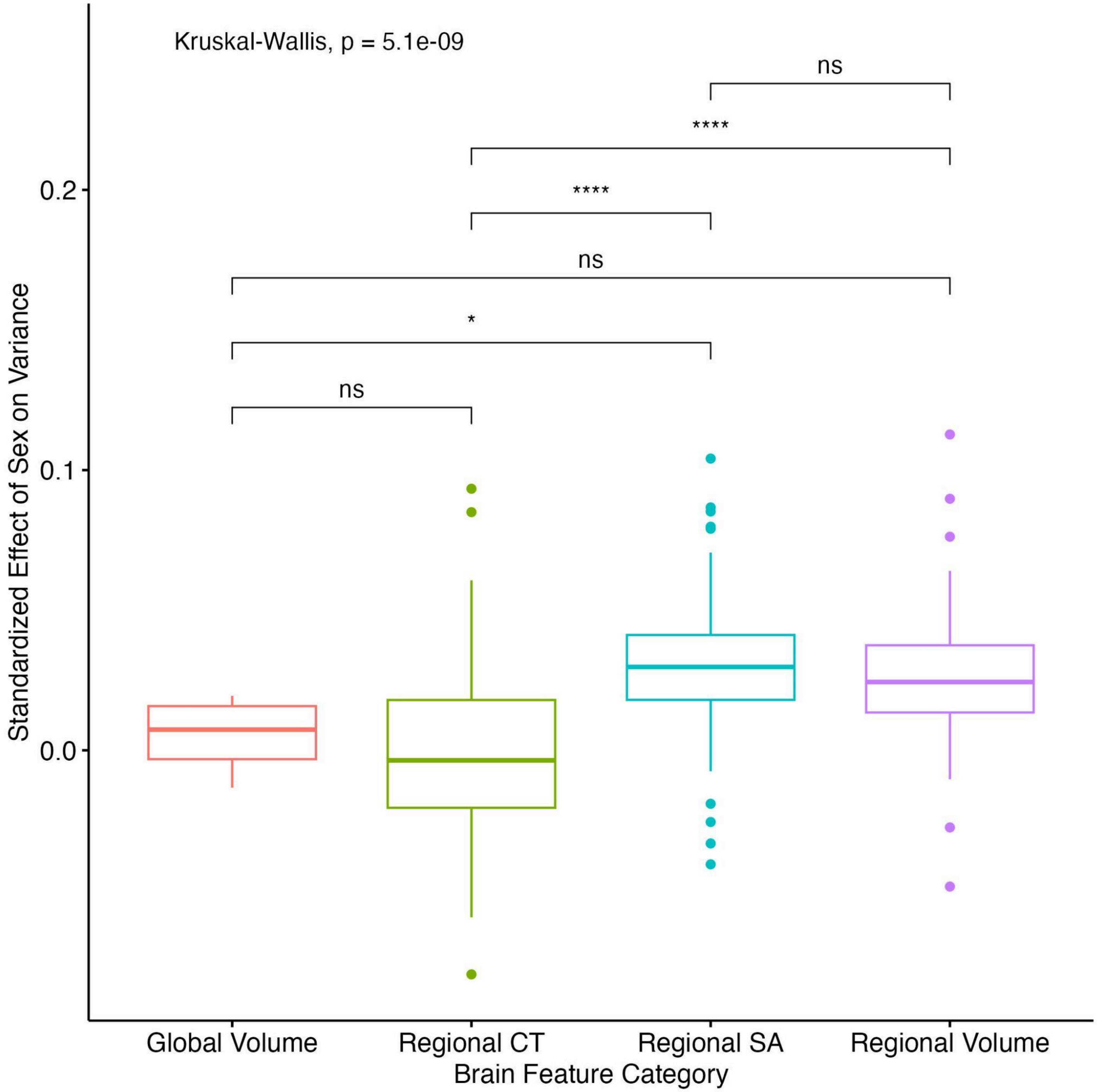

**Supplemental Figure 11. Sex has smaller effects on cortical thickness features' variances than other regional phenotypes.** Abbrv: CT, cortical thickness; SA, surface area; ***, $p < 0.001$; **, $p < 0.01$, FDR-corrected.

**Section III. Replication without Extreme Centiles**

To determine whether a small number of subjects with very high or very low centile scores drove differences across ComBat methods, we repeated our statistical comparisons after removing subjects with "extreme" centiles in a given feature. We defined extreme centiles as those >95% or <5% when calculated from raw, "unharmonized" data. First, we identified and removed extreme centiles across our 100 replications. We then compared the remaining centile errors across ComBat methods, using two-tailed t-tests of centile error ranks with Welch's correction, controlling FDR for 1248 comparisons within each replication. We also repeated our assessments of sex-differences in centile errors and ComBat-induced sex biases in centile displacement after taking the median of sex-differences within feature categories. Second, we applied these procedures to our 11 samples of synthetic sites with varying Male:Female ratios to assess whether extreme subjects drove differences in ComBat methods' performance across degrees of sex imbalance.

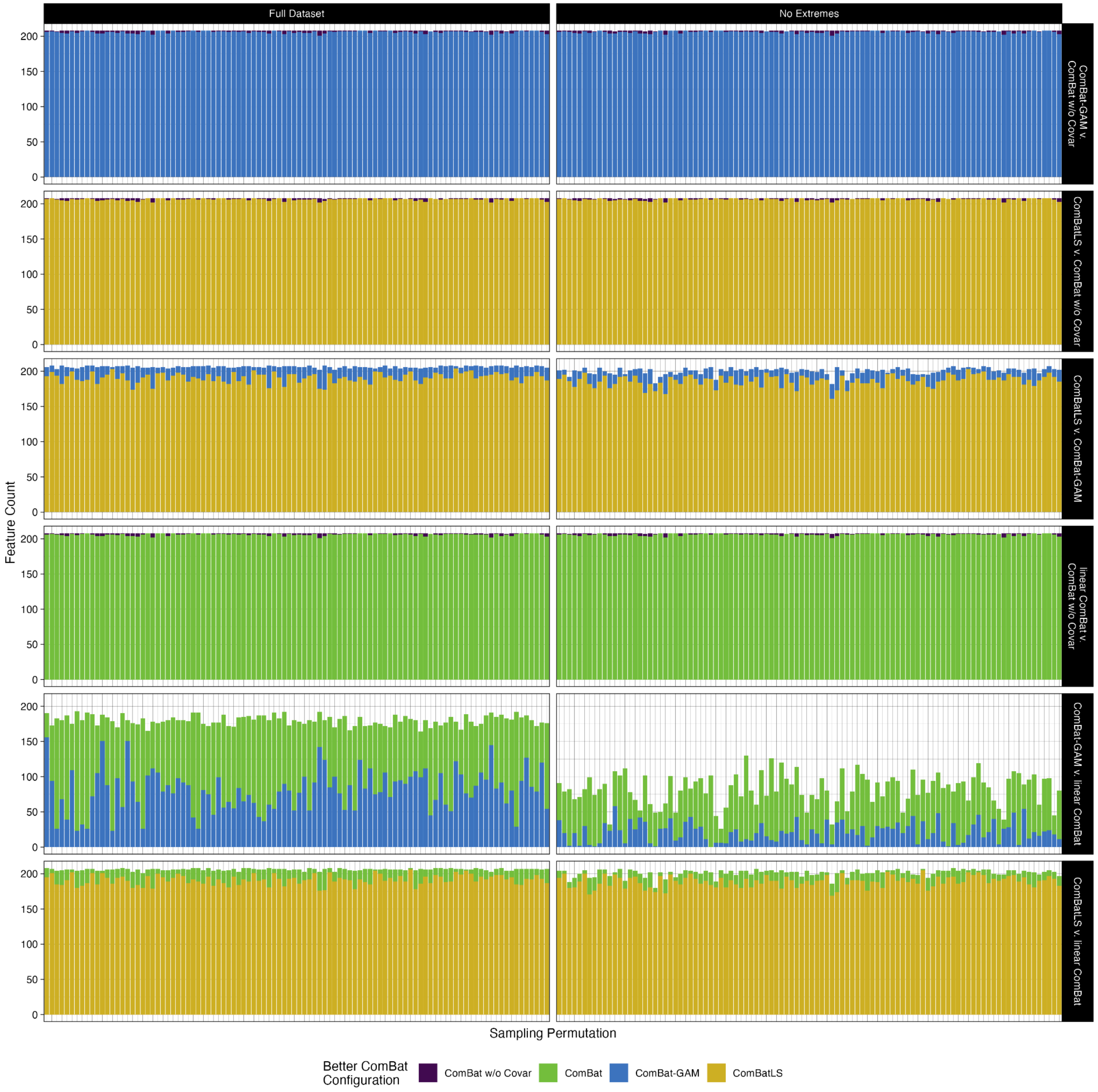

**Supplemental Figure 12. Extreme phenotypes do not drive differences in absolute centile errors between ComBat methods.** Comparison of pairwise tests of absolute centile errors between ComBat methods when centiles with raw values above 95% or below 5% are excluded. Absolute centile errors were compared within each brain featureacross 100 sampling permutations. Fill indicates the ComBat method that produces significantly smaller absolute centile errors, FDR-corrected across 1248 tests (208 features x 6 ComBat method pairings) within each permutation.

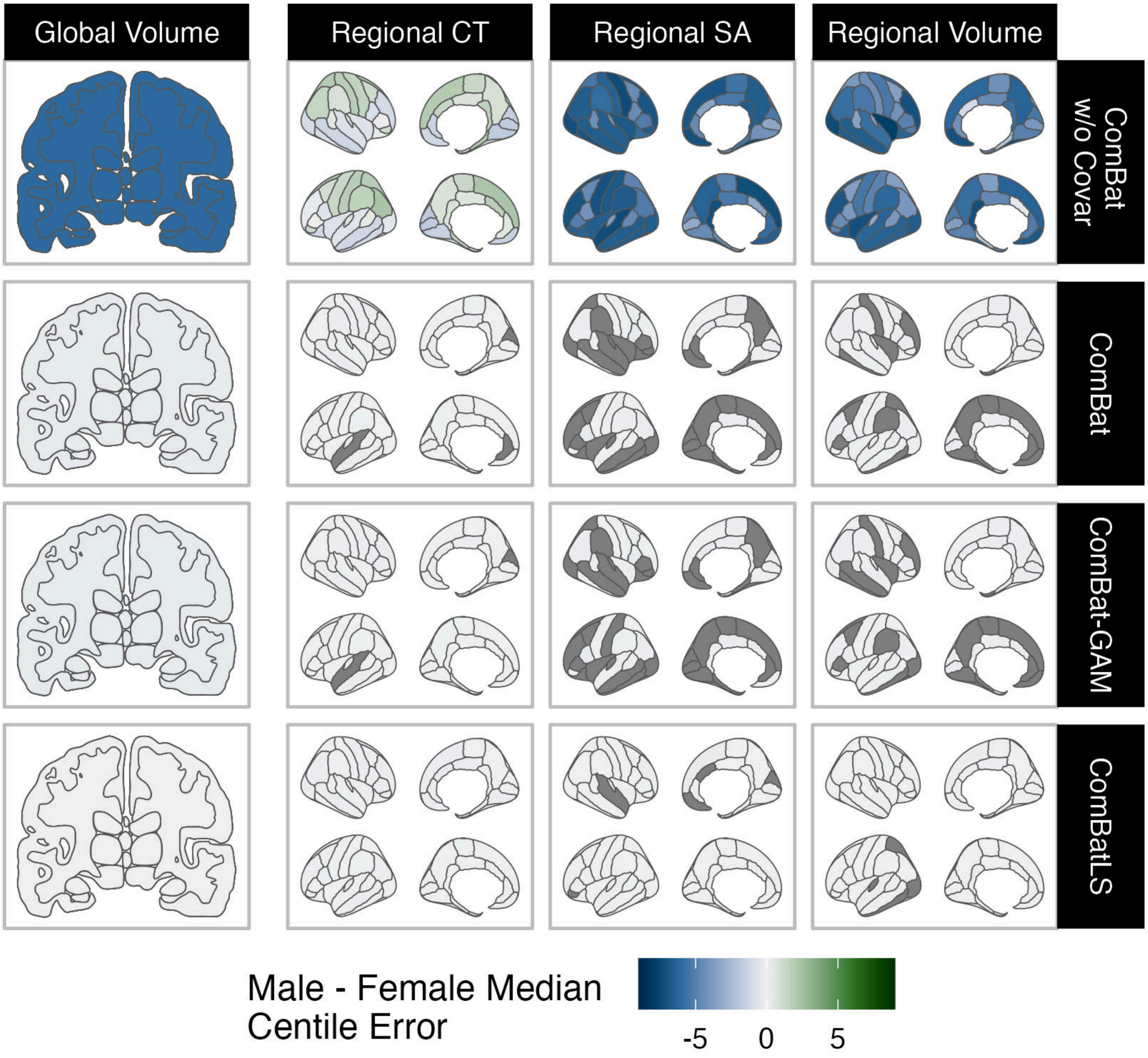


**Supplemental Figure 13. Significant differences in males' and females' median centile errors across brain features and ComBat methods when extreme features are excluded.** Positive centile errors (green) indicate that males' centiles tend to be overestimated relative to females'. Abbrv: CT, cortical thickness; SA, surface area.

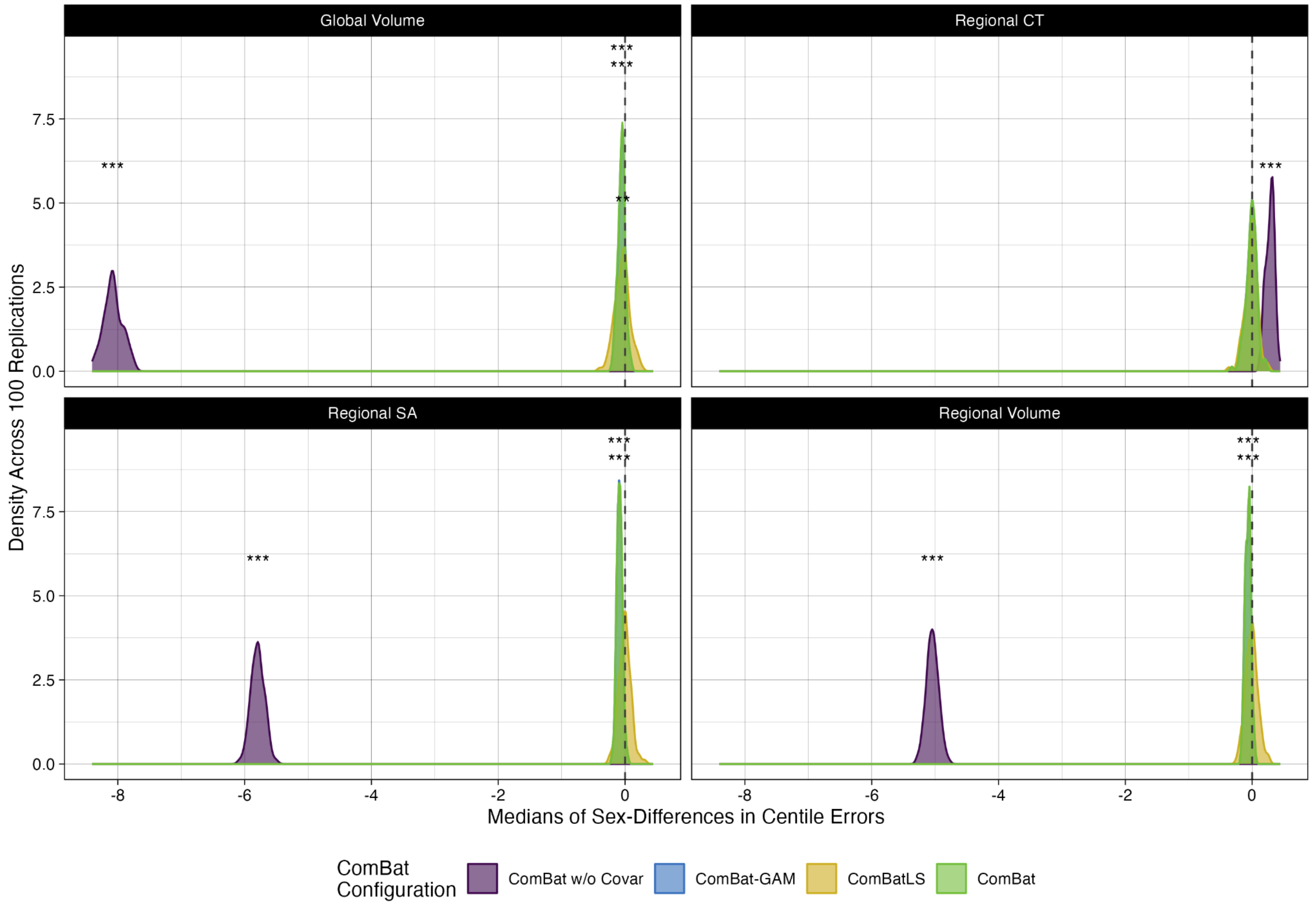


**Supplemental Figure 14. Density plots of median sex differences in centile errors induced by different ComBat methods across 100 replications when extreme phenotypes are excluded**. Abbrv: CT, cortical thickness; SA, surface area; ***, $p < 0.001$; **, $p < 0.01$, FDR-corrected.

### Section IV. Z-score analyses

Z-scores for each feature derived from centile scores using R's *qnorm()* function. To prevent infinite z-scores, centiles of 0 and 1 were estimated as 1e-25 and 0.99999999999999994, respectively. As with centile scores, analyses of z-scores were repeated without extreme scores, here defined as z-scores less than -2 or greater than 2.

#### A) Main Results

Pairwise comparisons of ComBat methods' absolute Z-score errors for 208 brain features across 100 replications

| Method Producing Smaller Absolute Errors | N Features | % Features |
|---|---|---|
| ComBat-GAM vs ComBat w/o Covariates | | |
| ComBat w/o Covariates | 191 | 0.92% |
| ComBat-GAM | 20600 | 99.04% |
| ComBatLS vs ComBat w/o Covariates | | |
| ComBat w/o Covariates | 156 | 0.75% |
| ComBatLS | 20634 | 99.20% |
| ComBatLS vs ComBat-GAM | | |
| ComBat-GAM | 1617 | 7.77% |
| ComBatLS | 18992 | 91.31% |
| Linear ComBat vs ComBat w/o Covariates | | |
| ComBat w/o Covariates | 189 | 0.91% |
| Linear ComBat | 20599 | 99.03% |
| Linear ComBat vs ComBat-GAM | | |
| ComBat-GAM | 7639 | 36.73% |
| Linear ComBat | 10398 | 49.99% |
| Linear ComBat vs ComBatLS | | |
| ComBatLS | 19030 | 91.49% |
| Linear ComBat | 1580 | 7.60% |

**Supplemental Table 2**. Pairwise tests of absolute z-score errors within each of 208 brain features replicated across 100 subject resamplings. All tests conducted as pairwise, two-tailed t-tests of ranks with Welch's correction. FDR-corrected across 1248 tests (208 features x 6 ComBat method pairings) within each sampling permutation.

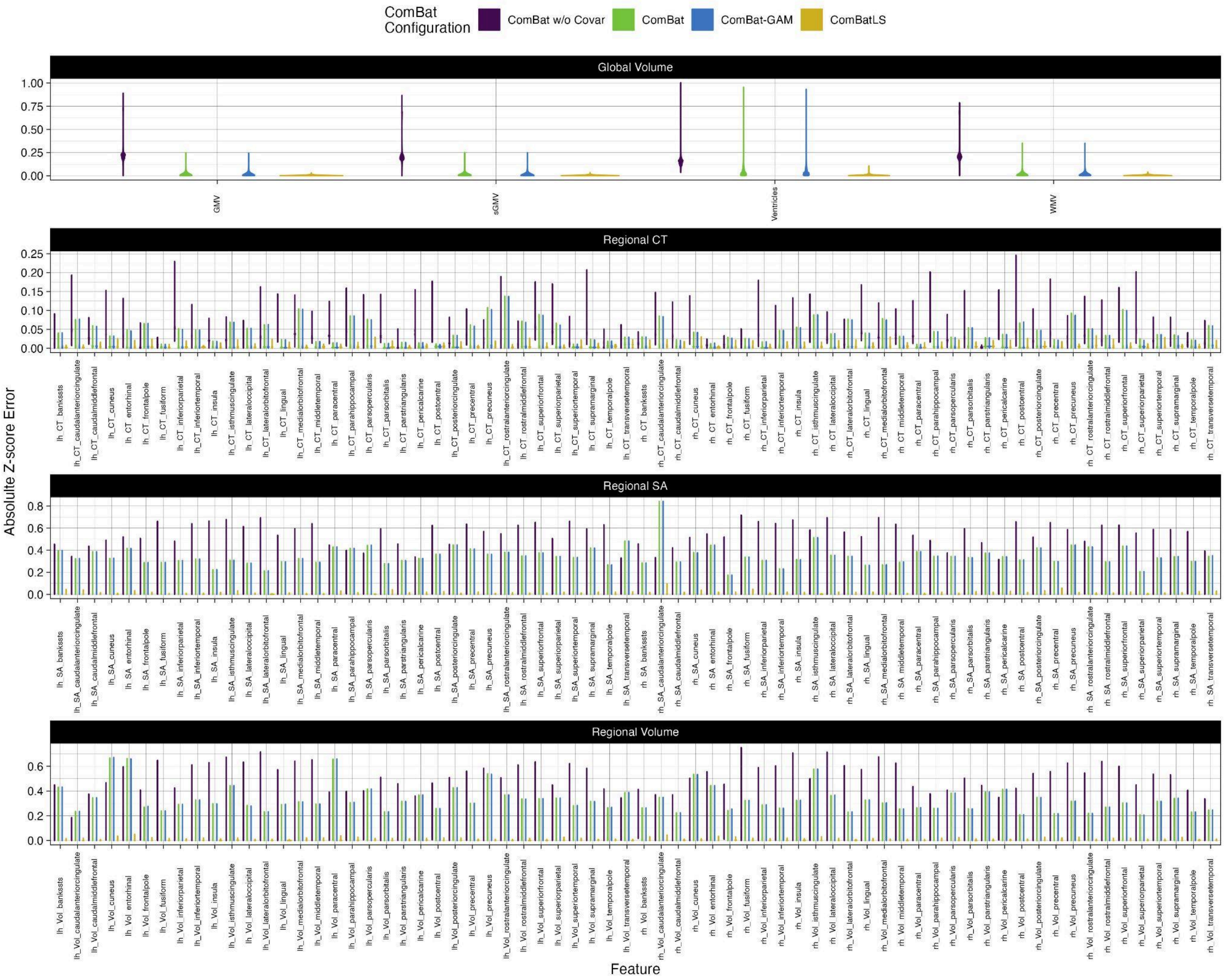


**Supplemental Figure 15. Absolute z-score errors across brain features and ComBat methods.** Violin plots of absolute z-score errors across 208 brain features.

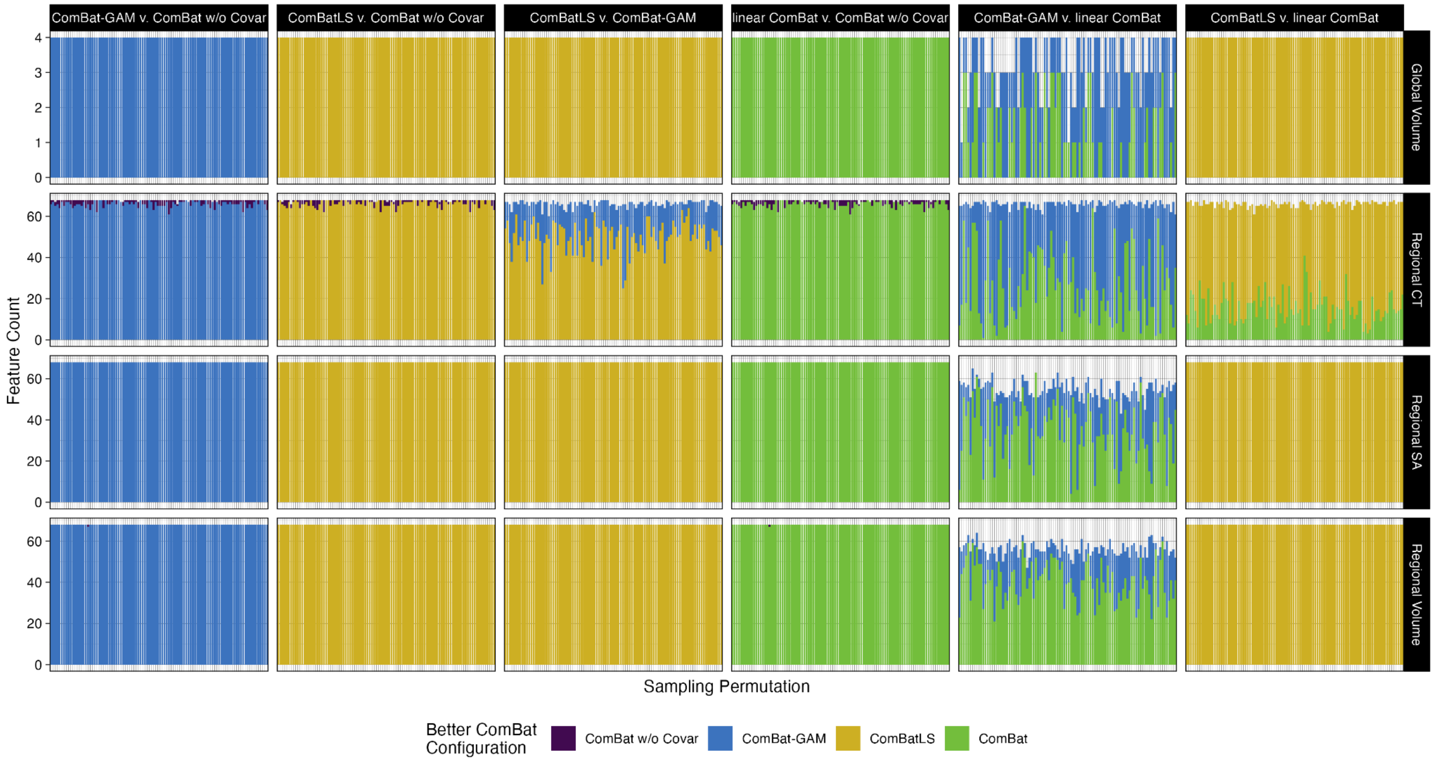


**Supplemental Figure 16. Pairwise comparisons of absolute z-score errors across ComBat methods within each brain feature across 100 sampling replications**. Fill indicates the ComBat method that produces significantly smaller absolute z-score errors, FDR-corrected across 1248 tests (208 features x 6 ComBat method pairings) within each permutation.

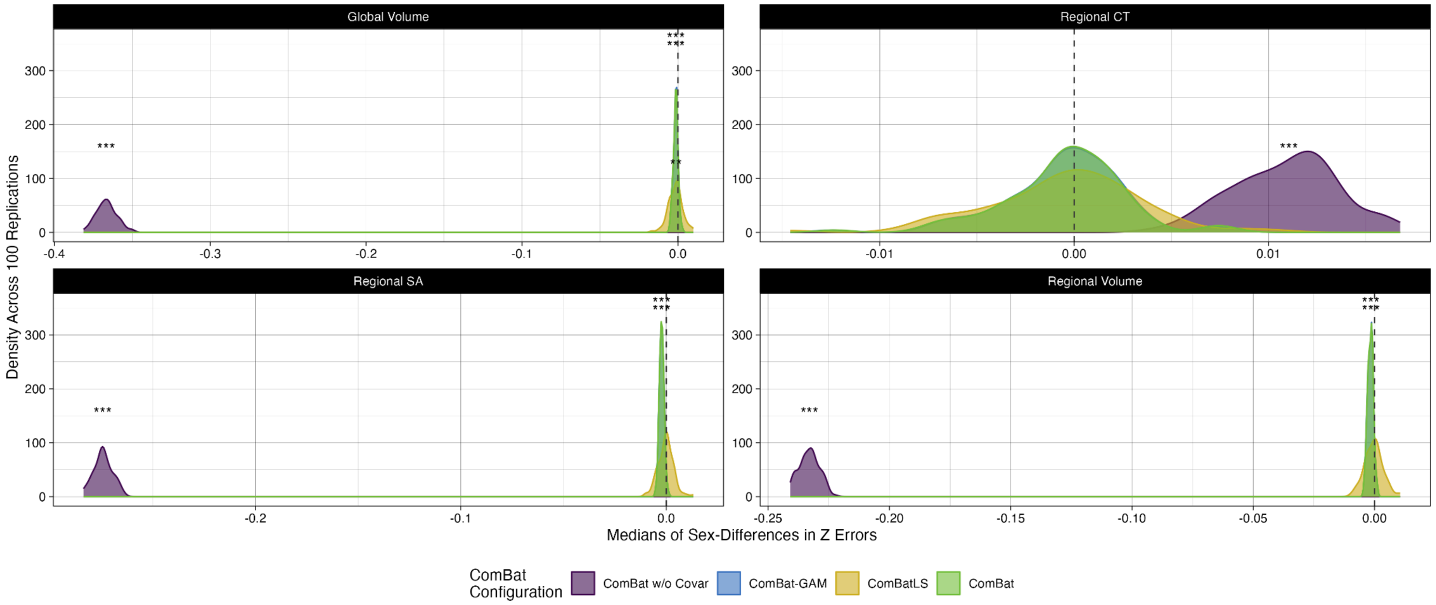


**Supplemental Figure 17. Density plots of median sex differences in z-score errors induced by different ComBat methods within phenotype categories across 100 replications.** Abbrv: CT, cortical thickness; SA, surface area; ***, $p < 0.001$; **, $p < 0.01$, FDR-corrected.

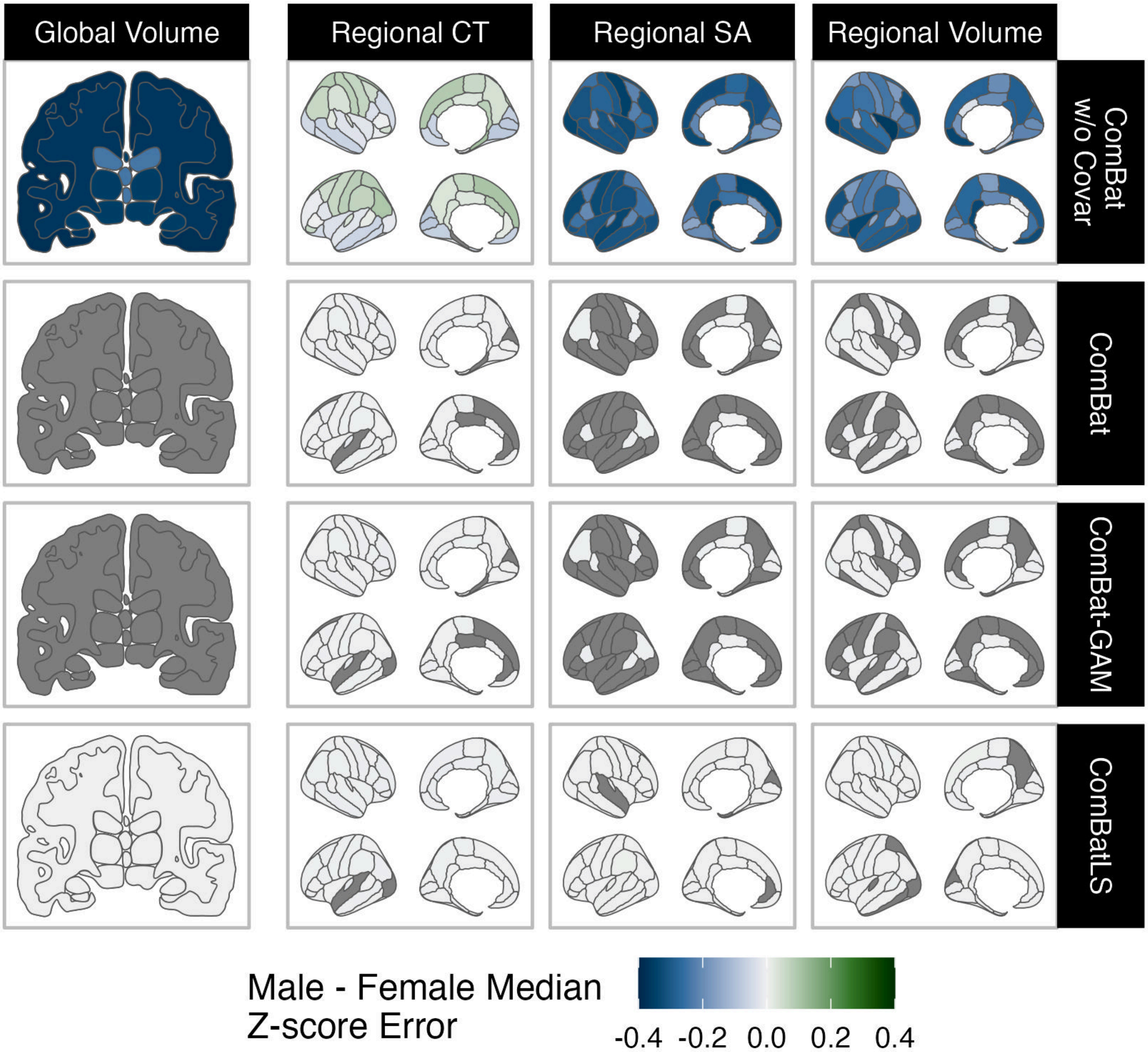


**Supplemental Figure 18. Significant differences in males' and females' median z-score errors across brain features and ComBat methods.** Positive centile errors (green) indicate that males' z-scores tend to be overestimated relative to females'. Abbrv: CT, cortical thickness; SA, surface area.

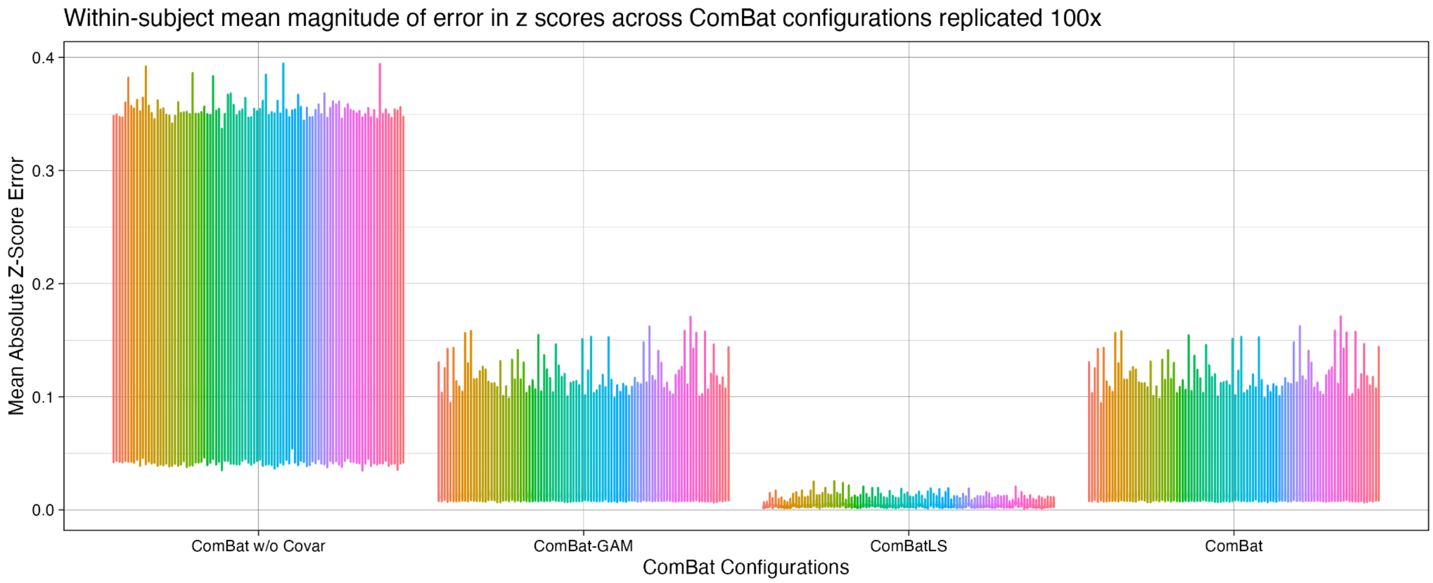


**Supplemental Figure 19. Subjects' mean absolute z-score error across ComBat configurations and 100 sampling replications.** Violin plots of absolute z-score error for 208 features averaged within subject. Fill corresponds to sampling replication.

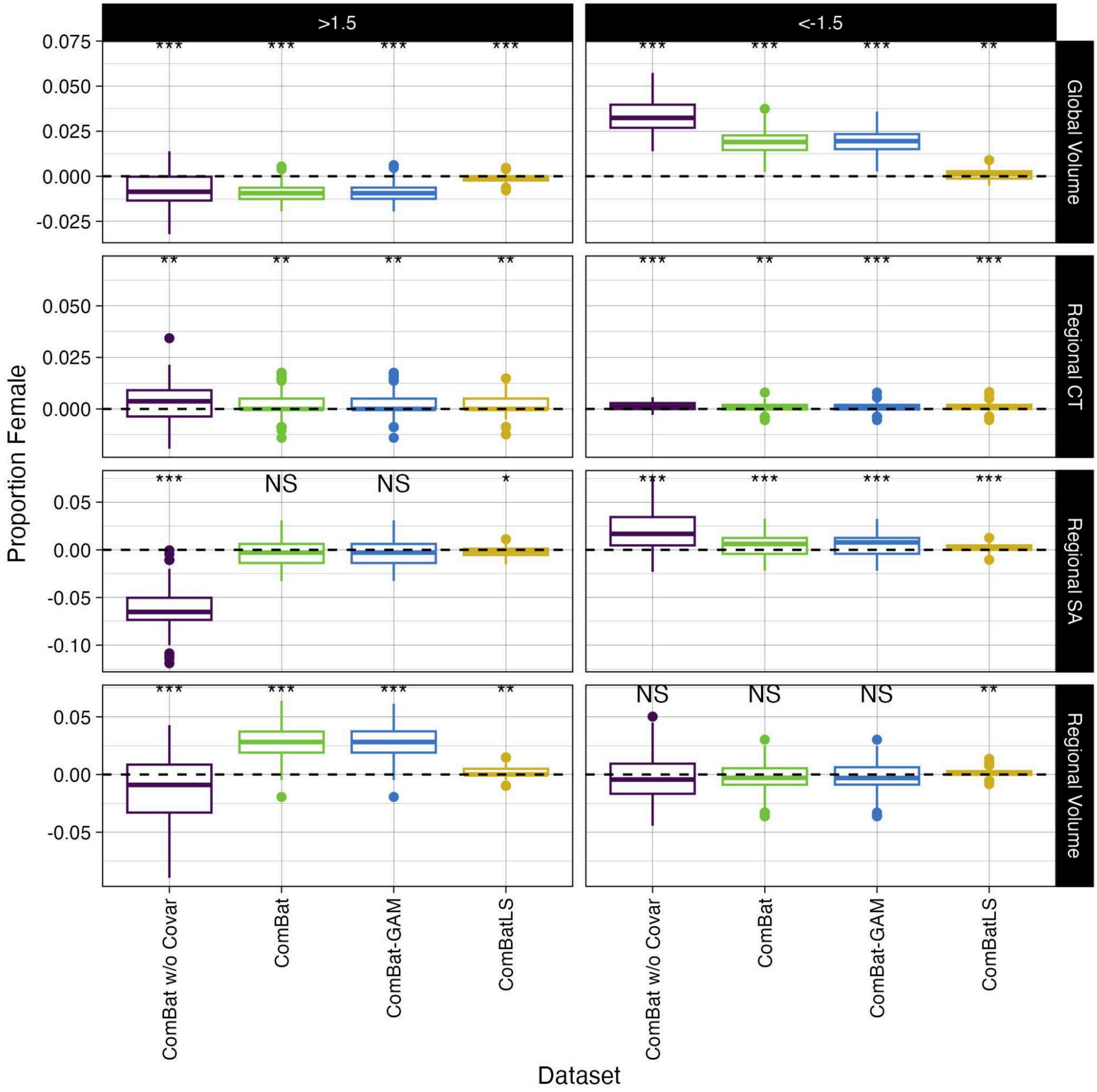


**Supplemental Figure 20. Over- or under-representation of females among individuals with extreme z-scores across ComBat methods.** Bias in the proportion of females with low (<-1.5) or high (>1.5) mean z-scores across 100 sampling replications. Positive values indicate a higher proportion of females than "true" mean z-scores calculated from unharmonized data (dashed line). Abbrv: ***, $p < 0.001$; **, $p < 0.01$, FDR-corrected.

### B) Varying M:F ratios

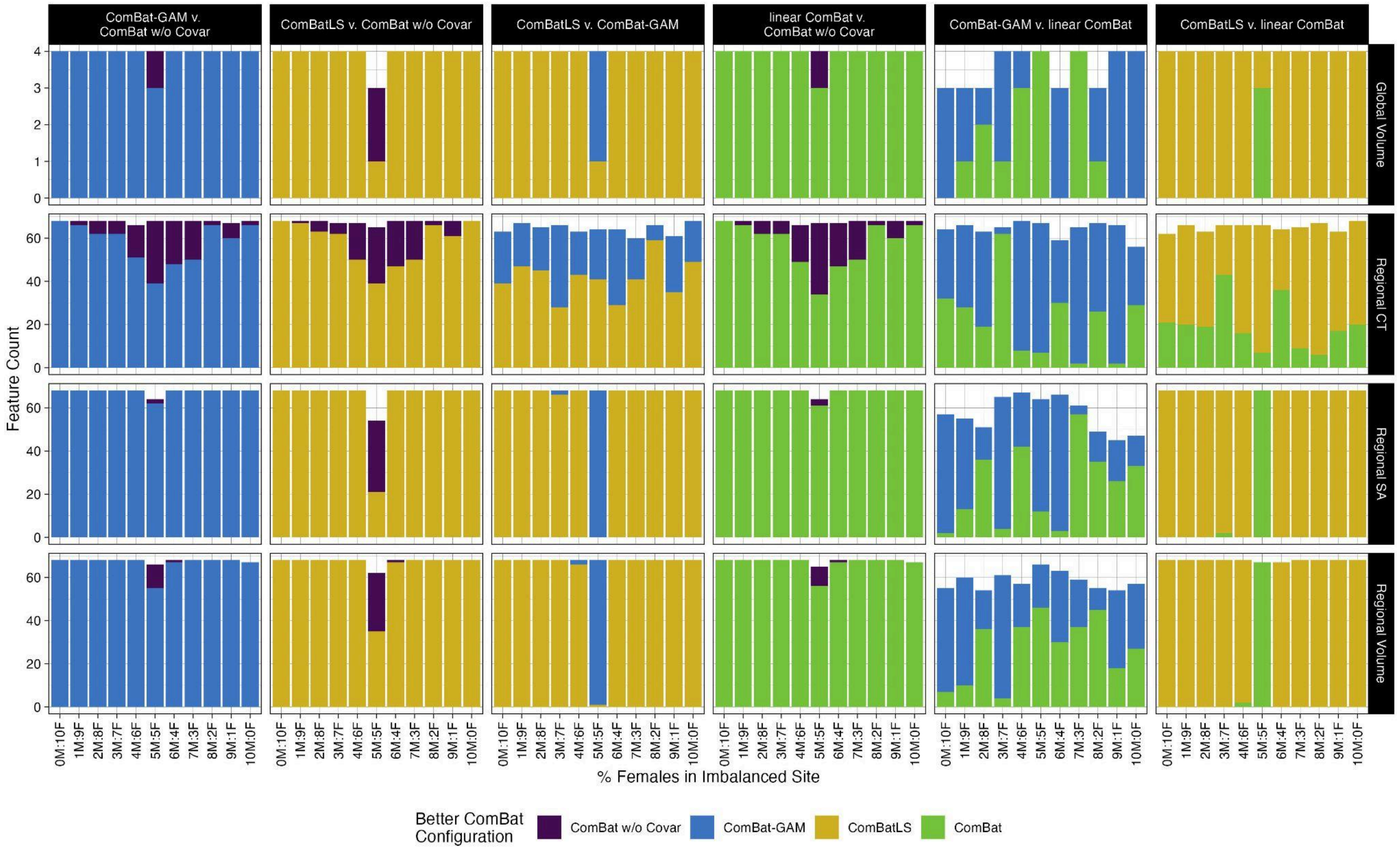


**Supplemental Figure 21. Magnitude z-score errors compared pairwise between ComBat methods across varying levels of sex-imbalances in simulated sites.** Fill indicates ComBat method with significantly lower absolute z-score errors for a given feature, FDR-corrected across simulated sex ratios. Abbrv: CT, cortical thickness; SA, surface area.

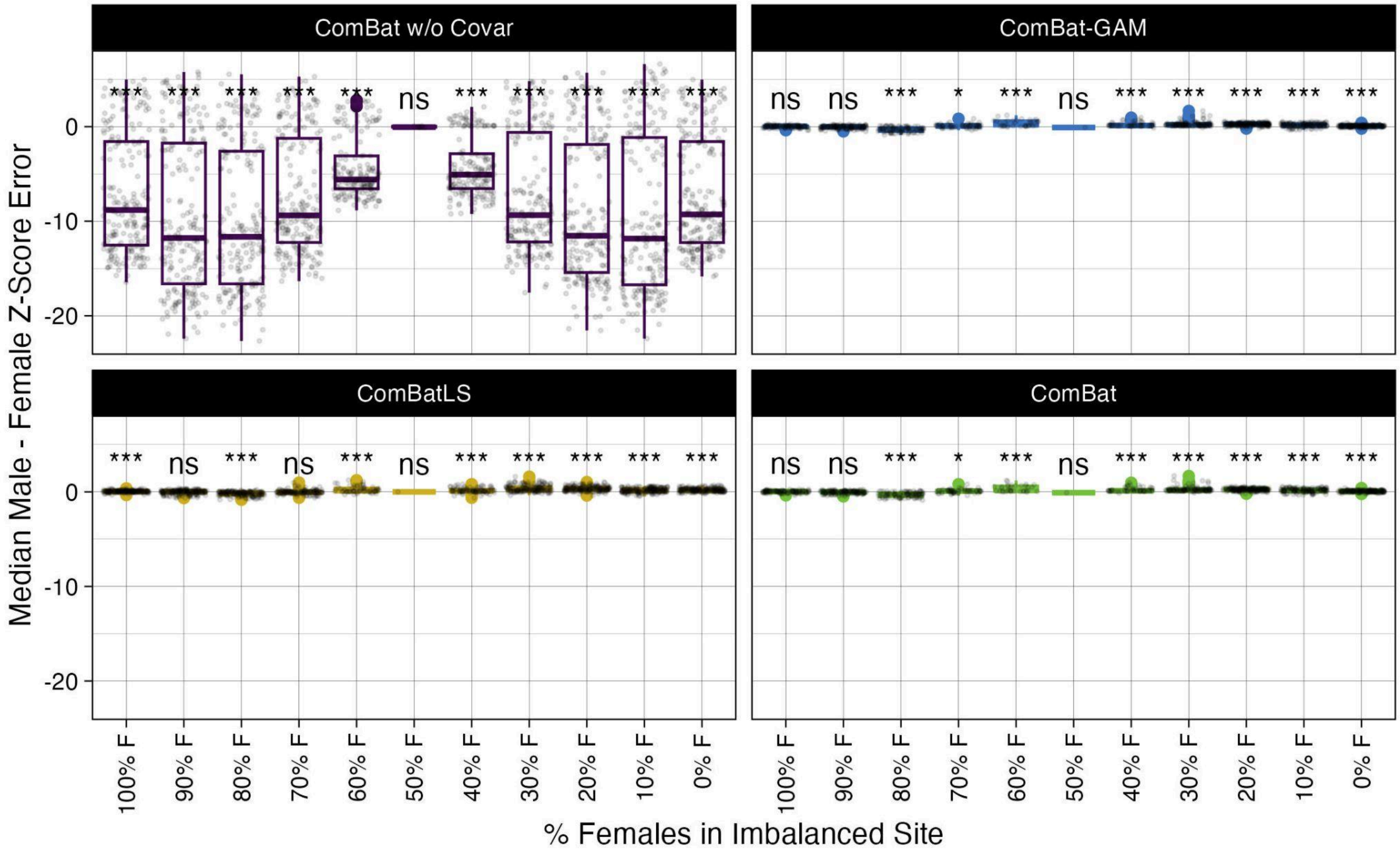


**Supplemental Figure 22. Sex-biases in z-score errors induced by various ComBat methods across varying degrees of sex-imbalance.** Points show brain features with significant differences in the distributions of males' and females' z-score errors (FDR corrected). Boxplots show median male - median female z-score errors across these features when centiles are derived from data harmonized by different ComBat methods. ComBat without covariate preservation induces strong biases wherein males' z-scores are underestimated relative to females', particularly as simulated sites become more imbalance for sex. Abbrv: ***, $p < 0.001$; **, $p < 0.01$, FDR-corrected.

**C) Without Extreme Z-scores**

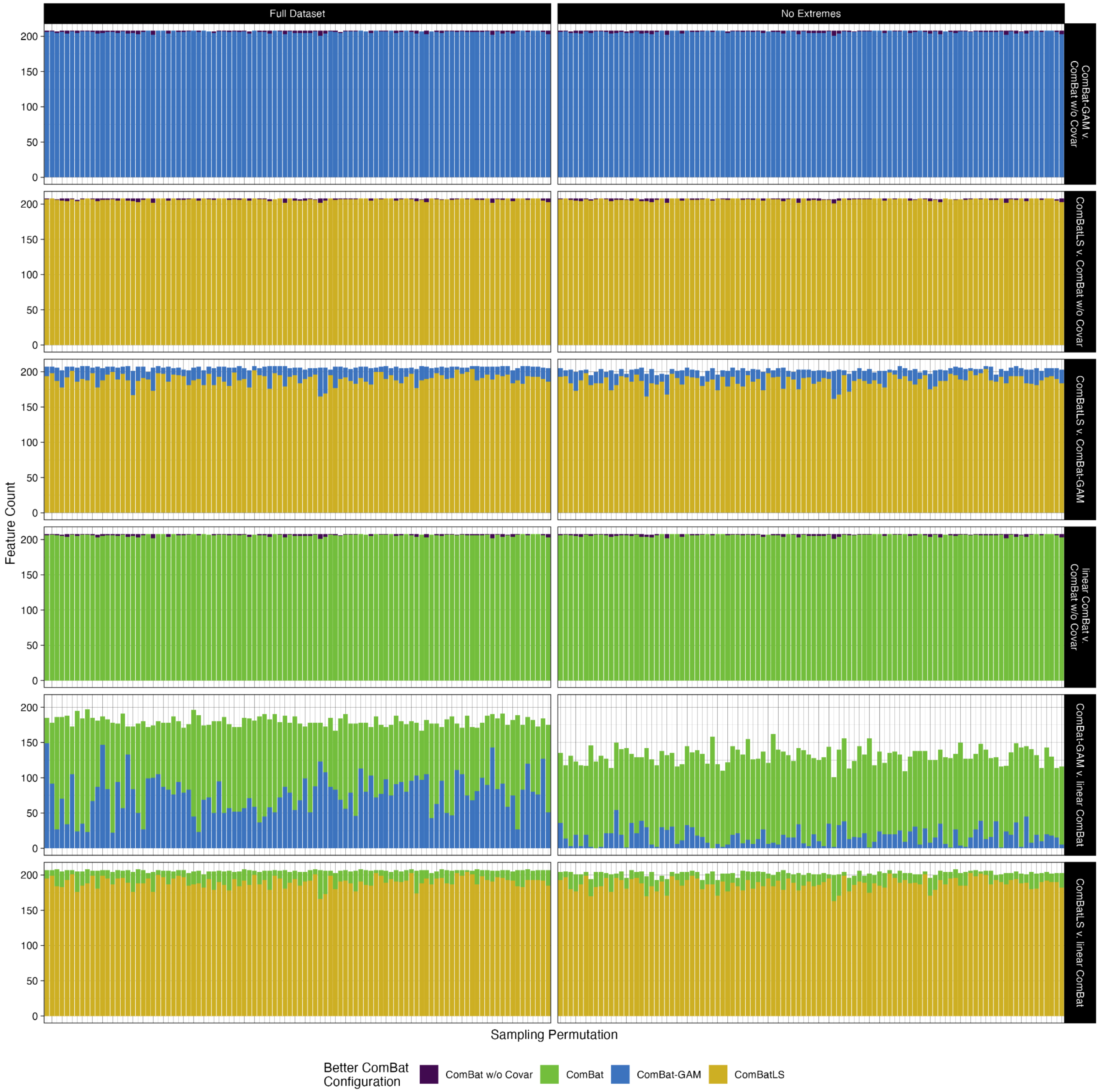


**Supplemental Figure 23. Extreme phenotypes do not drive differences in absolute z-score errors between ComBat methods.** Comparison of pairwise tests of absolute z-score errors between ComBat methods when z-scores with raw values above 2 or below -2 are excluded. Absolute z-score errors were compared within each brain feature across 100 sampling permutations. Fill indicates the ComBat method that produces significantly smaller absolute errors, FDR-corrected across 1248 tests (208 features x 6 ComBat method pairings) within each permutation.

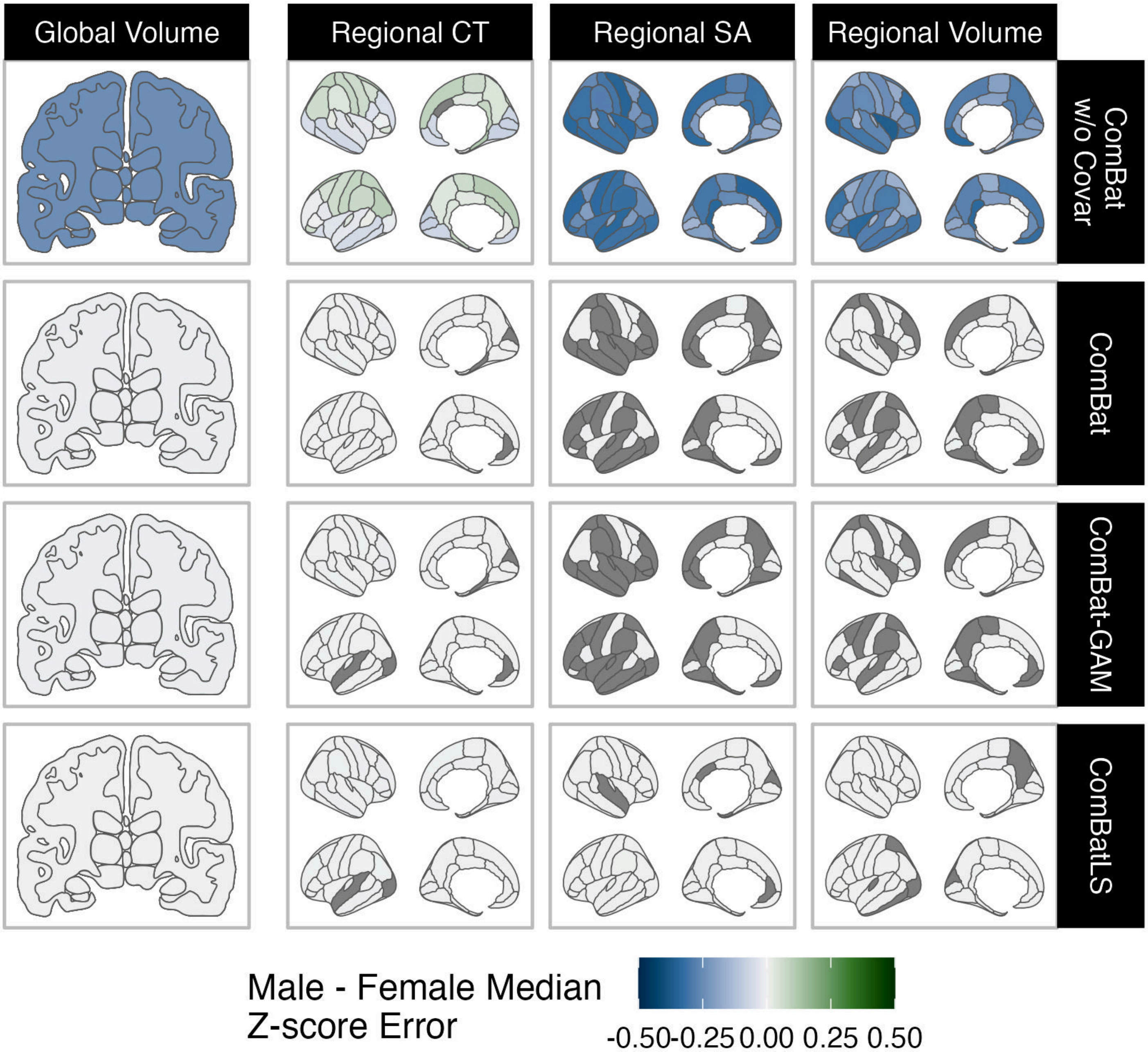


**Supplemental Figure 24. Significant differences in males' and females' median z-score errors across brain features and ComBat methods when extreme features are excluded.** Positive z-score errors (green) indicate that males' z-score tend to be overestimated relative to females'. Abbrv: CT, cortical thickness; SA, surface area.

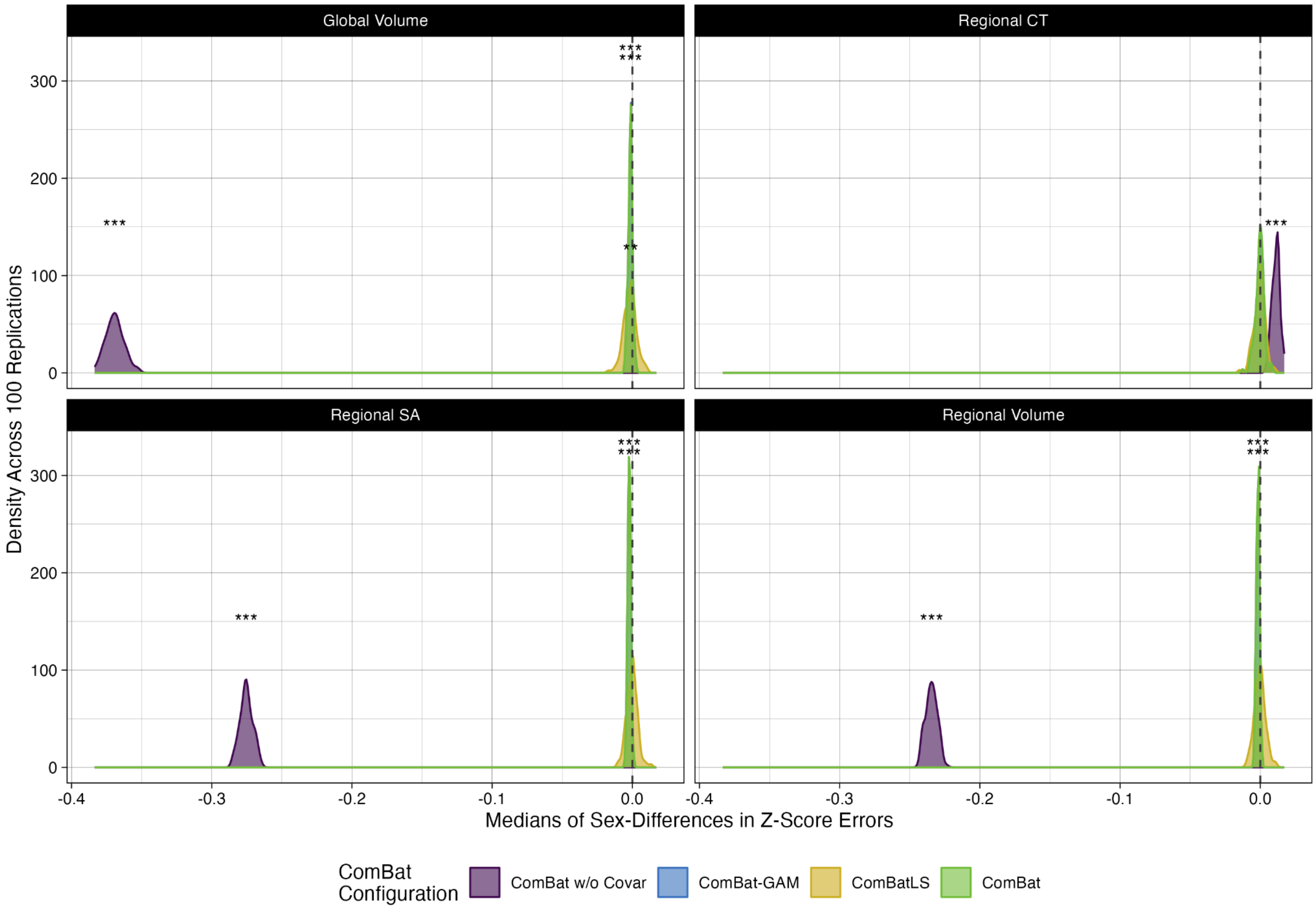


**Supplemental Figure 25. Density plots of median sex differences in z-score errors induced by different ComBat methods across 100 replications when extreme phenotypes are excluded**. Abbrv: CT, cortical thickness; SA, surface area; ***, $p < 0.001$; **, $p < 0.01$, FDR-corrected.

**<u>Section V. Comparisons of ComBatLS and ComBat-GAM for harmonizing consortium data</u>**

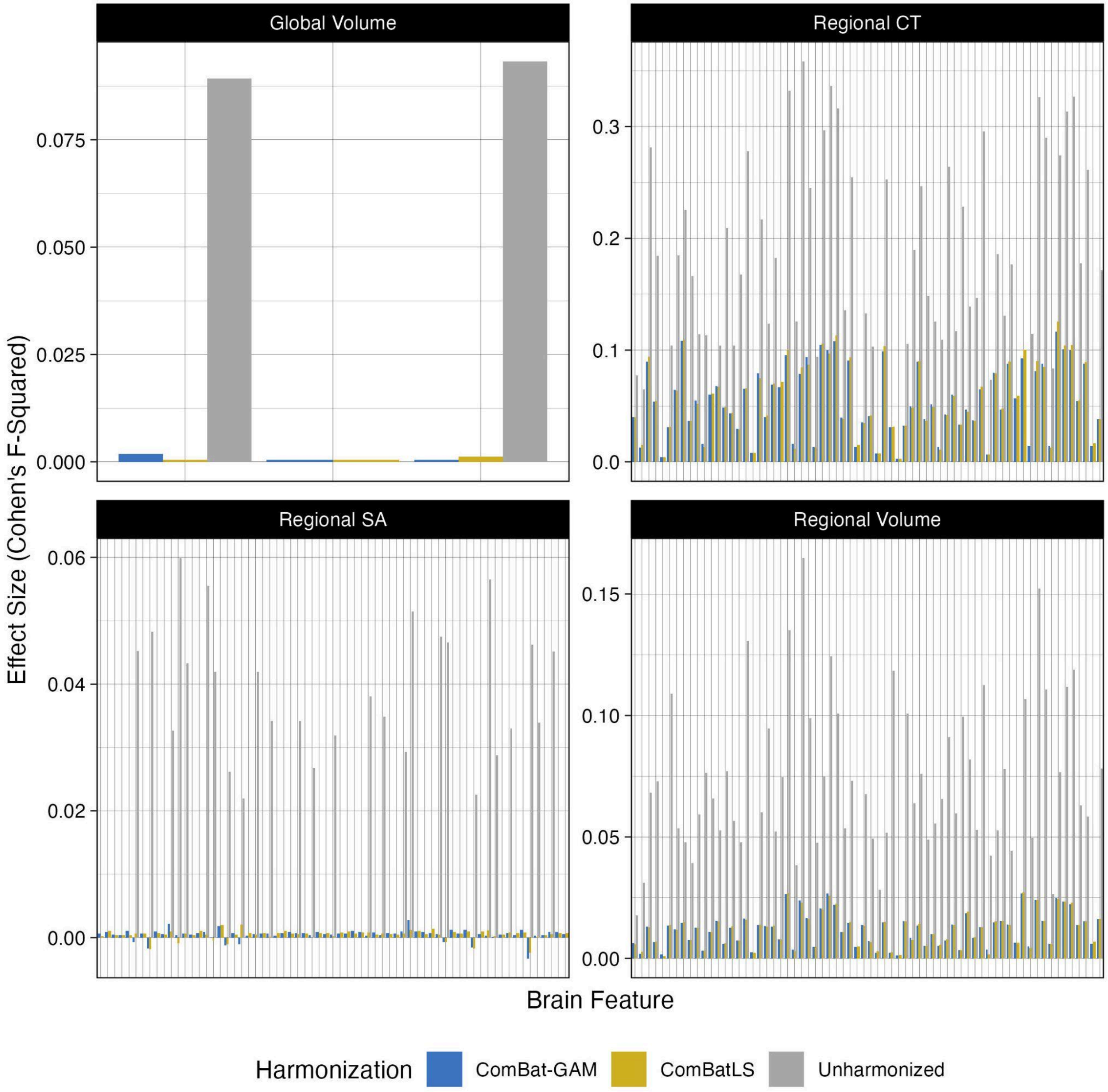


**Supplemental Figure 26. Residual effects of study following harmonization with ComBatLS or ComBat-GAM relative to unharmonized data.** Effect size for study in each brain feature's gamlss growth chart after harmonization. Note that not all brain feature's models converged in all datasets. Abbrv: CT, cortical thickness; SA, surface area.

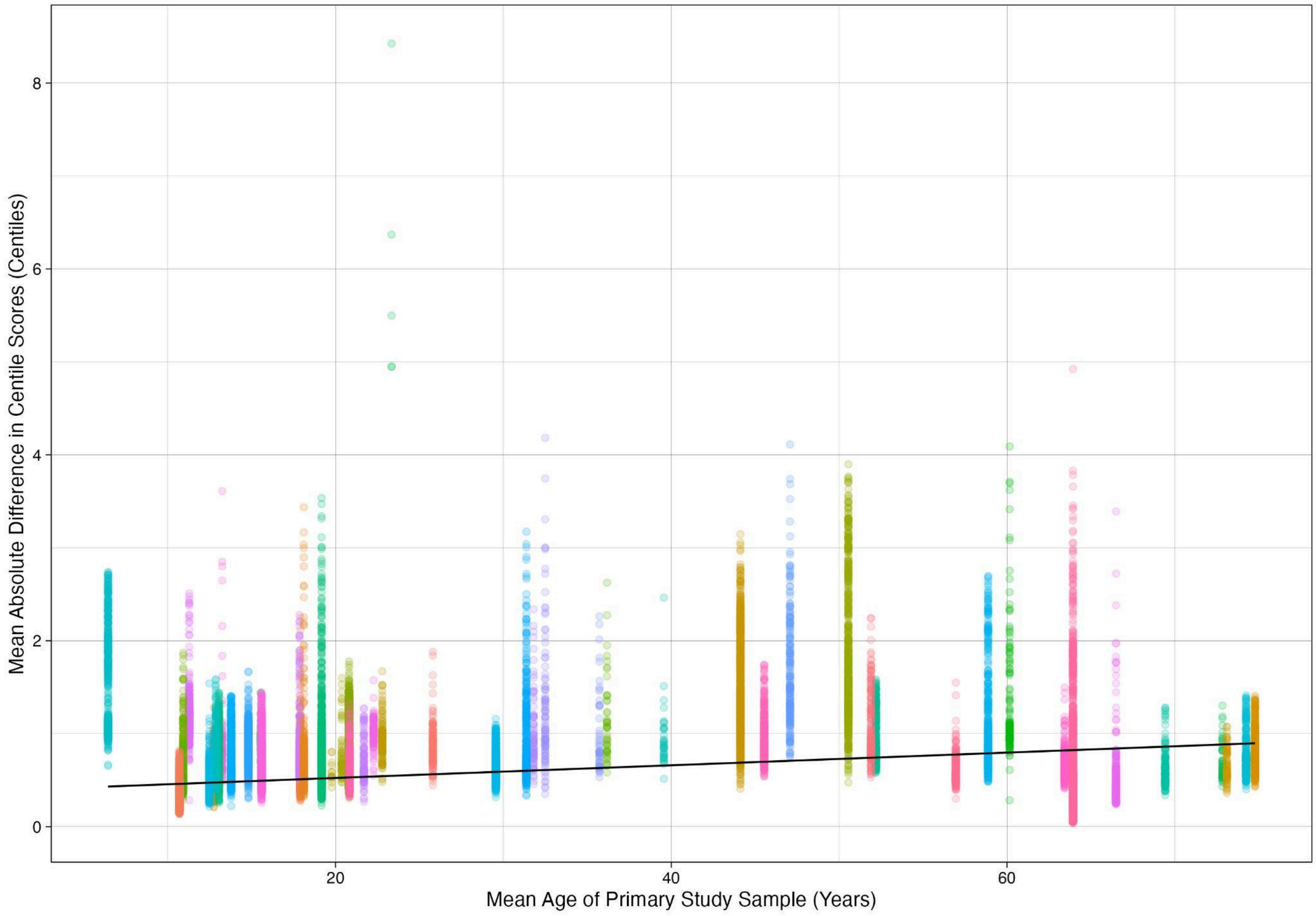


**Supplemental Figure 27. Absolute differences in ComBatLS and ComBat-GAM-derived centiles are related to study's mean age**. Y-axis shows mean absolute difference in a subject's centile scores across brain features when derived from ComBatLS- or ComBat-GAM-harmonized data. Each point represents one individual while fill represents primary study. Line shows marginal effect of mean study sample age, controlling for sample size. Gray band (not visible) represents 95% confidence interval of marginal association (Beta = 0.007 centiles, p < 2e-16).

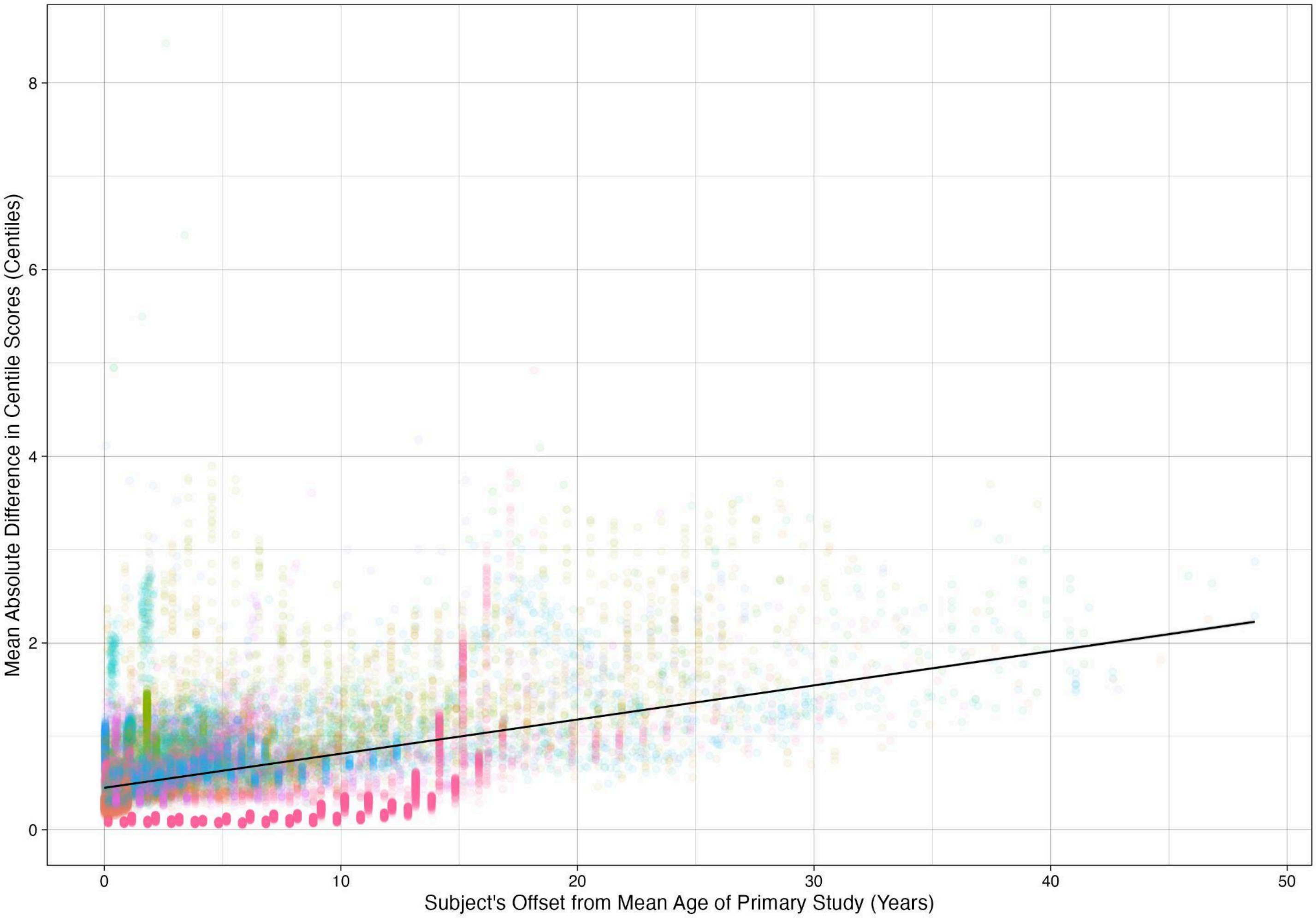


**Supplemental Figure 28. Absolute differences in ComBatLS and ComBat-GAM-derived centiles are larger in individuals whose ages are farther from their study's mean**. Y-axis shows mean absolute difference in a subject's centile scores across brain features when derived from ComBatLS- or ComBat-GAM-harmonized data. X-axis denotes the absolute difference in years between the individual subjects' age and the mean age of subjects in their primary study's sample. Each point represents one individual while fill represents primary study. Line shows marginal effect of absolute age difference, controlling for sample size. Gray band (not visible) represents 95% confidence interval of marginal association (Beta = 0.037 centiles, $p < 2e-16$).
